# Comparative Analysis of Tylosema esculentum Mitochondrial DNA Revealed Two Distinct Genome Structures

**DOI:** 10.1101/2023.03.27.534440

**Authors:** Jin Li, Christopher Cullis

## Abstract

*Tylosema esculentum* (marama bean), an underutilized legume with edible and nutritious seeds, has the potential to improve local food security in southern Africa. This study investigated the diversity of marama mitogenomes by mapping sequencing data from 84 individuals to the previously published reference mitogenome. Two distinct germplasms were found, and a new mitogenome structure containing three circular molecules and one long linear chromosome was identified, with a unique 2,108 bp fragment and primers were designed on that for marama mitogenome typing. This structural variation increases copy number of certain genes, including *nad9*, *rrns* and *rrn5*. The two mitogenomes also differed at 230 loci, with only one nonsynonymous substitution in *matR*. The evolutionary analysis suggested that the divergence of marama mitogenomes may be related to soil moisture level. Heteroplasmy in the marama mitogenome was concentrated at specific loci, including 127,684 bp to 127,686 bp on chromosome LS1 (OK638188), and may be crucial in the evolution. Additionally, the mitogenomes of marama contained a cpDNA insertion of over 9 kb with numerous polymorphisms, resulting in the loss of function of the genes on that segment. This comprehensive analysis of marama mitogenome diversity may provide valuable insight for future improvement of the bean.

**Highlight:** The analysis of 84 marama mitogenomes revealed two germplasms and the structural variation affects certain gene copy numbers. Soil moisture levels may have played important roles in the mitogenome divergence.

## Introduction

*Tylosema esculentum*, also known as marama bean and gemsbok bean, is a non-nodulated legume from the Fabaceae family, Cercidoideae subfamily (LPWG, 2017; Sinou et al., 2020). Marama is endemic to the Kalahari Desert and the surrounding arid and semi-arid regions of Botswana, Namibia, and South Africa (Bower et al., 1988). Marama has developed a special drought avoidance strategy by growing giant tubers that can weigh more than 500 pounds to store water, helping them survive the harsh conditions of long-term drought and little rainfall (Cullis et al., 2018). The seeds of marama are edible and nutritious, with a protein content of approximately 30-39% dry matter (dm) and a lipid content of 35-48% dm, comparable to those found in commercial crops soybean and peanuts, respectively (Amarteifio, 1998; Belitz et al., 2004; Holse et al., 2010). Marama also contains many micronutrients and phytochemicals that are beneficial to human health (Jackson et al., 2010; Khan, 2018; Omotayo and Aremu, 2021). However, marama is still mainly collected from wild plants, and the domestication of marama has long been thought to have the potential to improve local food security. Marama usually does not flower and produce seeds until two to four years after planting, making traditional breeding very inefficient (Cullis et al., 2019). Therefore, selecting improved marama individuals and exploring the underlying genetic diversity is considered of great significance for the improvement of the bean (Cullis et al., 2022).

Mitochondria in eukaryotic cells are generally considered to have originated from the endosymbiosis of alpha-proteobacteria, although a number of changes have occurred since then, including the loss of many genes and their transfer to the nuclear genome (Andersson et al., 2002). The mitochondrial genomes of animals and plants have been found to vary greatly. Animal mitogenomes are usually small, only about 16 kb in size, and contain 37 genes (Boore, 1999). Plant mitogenomes are commonly larger, with 50 to 60 genes and expanded intergenic non-coding regions that result from DNA transfer from other cellular compartments or even from different organisms (Unseld et al., 1997; Kazuyoshi and Kubo, 2010; Gualberto et al., 2014). In land plants, they range in size from 66 kb in *Viscum scurruloideum* to 11.3 Mb in *Silene conica* (Sloan et al., 2012; Skippington et al., 2015).

Plant mitochondrial genes are very conserved, are considered to play important roles in ATP synthesis, and are also related to plant fertility and environmental adaptation (Hanson, 1990; Budar and Roux, 2011; Heng et al., 2014). The base substitution rate of mitochondrial genes is lower than that of chloroplast genes, and it is far lower, only about one-tenth, of the rate observed in nuclear genes (Wolfe et al., 1987; Drouin et al., 2008). Plant mitogenomes evolve even up to 100 times slower than animal mitogenomes (Palmer and Herbon; 1988). However, the structure of plant mitochondrial genomes is very dynamic, with a large number of sequence rearrangements, and repeat-mediated homologous recombination plays an important role in its structural variations (Gualberto and Newton, 2017; Cole et al., 2018).

In the mitochondrial genomes of many angiosperms, repetitive sequences account for 5-10% of the total genome size, and in a few plants such as *S. conica* and *Nymphaea colorata*, the proportion can exceed 40% with repetitive fragments up to 80 kb in length (Sloan et al., 2012; Dong et al., 2018). Recombination mediated by short or intermediate length repeats is considered to be less frequent, but recombination on long repeats (>1 kb) is thought to occur more frequently and usually generates equimolar recombined molecules in the plant mitogenomes (Arrieta-Montiel, 2009; Guo et al., 2016; Li and Cullis, 2021).

The third-generation sequencing technology, such as PacBio, provides long reads with an average length of 10-25 kb, spanning the long repeats in the plant mitogenome. This makes the study of structural variations caused by the long repeat mediated recombination possible (Kozik et al., 2019; Hon et al., 2020). High sequencing coverage is also important, not only to make the genome assembly more reliable, but also to determine the proportion of different chromosome structures. It is also indispensable for the accurate assessment of nucleotide polymorphisms (Telenti et al., 2016). The latest PacBio HiFi sequencing with an extremely high accuracy of 99.9%, which needs little correction by the data generated from other sequencing platforms, further promotes the genome assembly (Miga et al., 2020; Naish et al., 2021).

Although plant mitochondrial genomes are often reported as one master circular chromosome, in reality, they often exist as multipartite structures. This includes a combination of linear molecules, branched molecules, and subgenomic circular molecules (Oldenburg and Bendich, 1996; Manchekar et al., 2006). For example, the mitogenome assembly of *Solanum tuberosum* was found to contain at least three autonomous chromosomes, including two small circular molecules and a long linear chromosome of 312,491 bp in length (Varré et al., 2019).

In the previously published study on the mitogenome of *T. esculentum*, two different equimolar structures were found to coexist in the same individual. There are two autonomous rings with a total length of 399,572 bp and five smaller circular molecules (Li and Cullis, 2021). These two structures are believed to be interchangeable by recombination on three pairs of long direct repeats (3-5 kb in length). As described in the study of *Brassica campestris* mitogenome, recombination on a pair of 2 kb repeats was postulated to split the 218 kb master chromosome into two subgenomic circular molecules of 135 kb and 83 kb in length (Palmer and Shields, 1984).

The previous comparative analysis of 84 *T. esculentum* chloroplast genomes has found two distinct germplasms. The two types of chloroplast genomes are different from each other at 122 loci and at a 230 bp inversion (Li and Cullis, 2023). Among many of these loci, heteroplasmy, the existence of two or more different alleles, could be seen, albeit one generally at a frequency below 2%. The occasional paternal leakage is considered to cause this phenomenon (Kvist et al., 2003; Luo et al., 2018). The reason for its stable inheritance is thought to be related to the developmental genetic bottleneck, but the specific mechanism is still unclear (Floros et al., 2018).

Research on heteroplasmy should avoid interference from homologous sequences of mitochondrial DNA in other organelles. In fact, the horizontal gene transfer of DNA from chloroplast genome to mitogenome and nuclear genome, or between mitogenome and nuclear genome are very common (Woloszynska et al., 2004; Keeling and Palmer, 2008). The transfer of DNA from mitogenome to chloroplast genome is very rare, but it has been reported in a few studies (Goremykin et al., 2008; Straub et al., 2013). Many chloroplast genes have been found to become pseudogenes and lost function after being inserted into the mitogenome, but the reason behind it is still unclear (Li et al., 2022).

Although the mechanisms underlying the effects of cytoplasmic activities on agronomic traits are not well understood, previous studies on potato cytoplasmic diversity have found that cytoplasmic types are directly related to traits such as tuber yield, tuber starch content, disease resistance, and cytoplasmic male sterility (Sanetomo and Gebhardt, 2015; Anisimova and Gavrilenko, 2017; Smyda-Dajmund et al., 2020). Furthermore, certain mtDNA types and certain chloroplast DNA types were found to be linked (Lössl et al., 1999). A comparative genomics analysis based on the chloroplast genomes of 3,018 modern domesticated rice cultivars found that their genotypes fall into two distinct clades, suggesting that the domestication of these cultivars may have followed two distinct evolutionary paths (Moner et al., 2020). These studies have the potential to reveal important selections occurring in organelle genomes, improve understanding of plant adaptation to different environments, and provide a basis for crop breeding to increase yield in corresponding environments.

The chloroplast genome is widely used in research on plant evolution, but the comparative analysis based on plant mitochondrial genome is not so extensive, and there is even less research on the mitochondrial diversity within the same species (Nikiforova et al., 2013; Carbonell-Caballero et al., 2015). In this study, the diversity of marama mitochondrial genome was analyzed by mapping the WGS reads of 84 *T. esculentum* individuals to the previously assembled reference mitogenome aiming to: (1) discover possible mitogenome structural diversity and the impact of structural variations on gene sequence and copy numbers; (2) compare the differential loci in mitogenomes of 43 independent marama individuals collected from different geographical locations in Namibia and South Africa to explore the divergences that have occurred and possible decisive environmental factors behind them; (3) look at heteroplasmy, the co-existence of multiple types of mitogenomes within the same individuals, and compare allele frequencies in related individuals to better understand the underlying cytoplasmic inheritance; (4) track polymorphisms accumulated in the chloroplast DNA insertion and interpret the fate of the inserted gene residues. In addition, the conserved protein-coding genes from the mitogenomes of *T. esculentum* and other Fabaceae species were compared to explore the evolutionary relationship between them.

## Materials and Methods

### Plant materials and DNA extraction

Samples 4 and 32 were two individuals in the greenhouse of Case Western Reserve University grown from seeds collected in Namibia at undocumented locations. They were identified by PCR amplification to have type 2 and type 1 germplasms, respectively. 1 g fresh young leaves were collected from the two plants and ground thoroughly with a pestle in a mortar containing liquid nitrogen. DNA was then extracted using a Quick-DNA HMW MagBead kit (Zymo Research) following the protocol. The double-stranded DNA was quantified by the Invitrogen^TM^ Qubit^TM^ 3.0 Fluorometer after mixing 5 μl DNA with 195 μl working solution, and 200 ng DNA was electrophoresed on a 1.5% agarose TBE gel at 40 V for 24 hours. The plant materials included another 84 marama individuals, 44 of which were plants grown in different geographical locations in Namibia and South Africa, and the remaining 40 were progeny plants grown from seeds collected there, as described in the previous study (Table S1) (Li and Cullis, 2023). The WGS Illumina reads of these 84 individuals are available and stored in the NCBI SRA database (PRJNA779273).

### High-throughput sequencing

DNA extracted from Samples 4 and 32 (10 micrograms or more per sample) was sent to the Genomics Core Facility at the Icahn School of Medicine at Mount Sinai for sequencing. The HiFi sequencing libraries were prepared using the SMRTbell® express template prep kit 2.0 (Pacific Biosciences, Menlo Park, CA, USA). SMRT sequencing was performed on four 8M SMRT® Cells (two per sample) on the Sequel® II system. 2,184,632 PacBio HiFi reads with a total length of 21.6 G bases were generated for Sample 4 and 498 Mb reads for Sample 32. In addition, whole-genome sequencing of DNA from fresh young leaves or embryonic axis of germinating seeds from the 84 samples collected in Namibia and South Africa was performed using the Illumina platform, as described in the previous study (Li and Cullis, 2023).

### Mitogenome assembly and annotation

The PacBio HiFi long reads were assembled using the HiCanu assembler (Nurk et al., 2020; https://canu.readthedocs.io/en/latest/quick-start.html#assembling-pacbio-hifi-with-hicanu). The input genome size was set to 2 Mb to obtain more complete organelle genome contigs. The PacBio long reads spanning the ends of the contigs were used to further scaffold the assembly. The assembled mitogenome was annotated using MITOFY (Alverson et al., 2010; https://dogma.ccbb.utexas.edu/mitofy/), BLAST (Johnson et al., 2008; https://blast.ncbi.nlm.nih.gov/Blast.cgi), and tRNAscan-se 2.0 (Chan et al., 2021; http://lowelab.ucsc.edu/tRNAscan-SE/). The assembly was verified by mapping Illumina reads and PacBio HiFi reads from different individuals to the generated mitogenome sequences using Bowtie 2 v2.4.4 (Langmead and Salzberg, 2012; https://github.com/BenLangmead/bowtie2) and pbmm2 v1.10.0 (https://github.com/PacificBiosciences/pbmm2), respectively. The alignment was visualized and checked in IGV (Robinson et al., 2011; https://software.broadinstitute.org/software/igv/), showing no ambiguity. The mitochondrial gene arrangement maps of *T. esculentum* were drawn by OGDRAW (Greiner et al., 2019; https://chlorobox.mpimp-golm.mpg.de/OGDraw.html).

### Mitogenome polymorphism

The program NUCmer in MUMmer4 (Marçais et al., 2018; https://mummer4.github.io/index.html) was used to locate highly conserved regions in the two types of mitochondrial genomes of *T. esculentum*. The alignment was visualized via a synteny plot drawn by the RIdeogram (Hao et al., 2020; https://github.com/TickingClock1992/RIdeogram) package in R.

The 2,108 bp type 2 unique fragment was blasted in the NCBI database for potential origin. Two pairs of primers were designed for mitogenome typing, using NCBI Primer-BLAST (Ye et al., 2012; https://www.ncbi.nlm.nih.gov/tools/primer-blast/), to amplify across the two ends of this fragment by PCR. The primers to amplify across the left end included the left forward primer (GAGACCGAGCGCAAGAACTA) and the left reverse primer (TCAGATGGCTAAACAGGCGG), and the product size was 990 bp. The primers to amplify across the right end included the right forward primer (CGCTCGTGACTCATTGAGGA) and the right reverse primer (TTGGTAAGCGGATGCTCTGG), and the product size was 289 bp. 20 uL of mixtures were prepared by separately mixing DNA from six randomly selected *T. esculentum* samples and Promega GoTaq Green Master Mix. The amplifications started with denaturation at 95 °C for 5 min, followed by 30 cycles of 95 °C for 45 s, 54 °C for 45 s, and 72 ° C for 1 min, and a final 72 °C for 5 min.

The Illumina reads from the 84 individuals were mapped to the type 1 reference mitogenome of *T. esculentum* (OK638188 and OK638189) using Bowtie 2 v2.4.4. The alignments were searched for SNPs and indels using SAMtools 1.7 mpileup (Li, 2011; http://www.htslib.org/) and BCFtools 1.8 call (Li, 2011; http://www.htslib.org/doc/1.8/bcftools.html), and visualized in IGV to find heteroplasmy manually. To minimize the interference of sequencing errors and strand bias, only alleles with a frequency of at least 2%, a Phred score above 20, and presence in strands in both orientations were recorded. Alleles in the mtDNA homologous reads in the nuclear genome were excluded. The alleles from the mitochondrial or chloroplast genomes were distinguished by their frequency at the differential loci on the 9,798 bp chloroplast DNA insertion.

### Mitogenome divergence

A pairwise comparison was performed on the divergent mitogenomes of *T. esculentum* (OK638188 and OK638189) and six other Fabaceae species, including *Cercis canadensis* (MN017226.1), *Lotus japonicus* (NC_016743.2), *Medicago sativa* (ON782580.1), *Millettia pinnata* (NC_016742.1), *Glycine max* (NC_020455.1), and *Vigna radiata* (NC_015121.1) using the program PROmer (Delcher et al., 2002) in MUMmer4 to detect the syntenic regions. The alignments were then visualized by the RIdeogram package in R. A synteny block diagram between these seven mitogenomes was drawn by Mauve 2.4.0 (Darling et al., 2004; https://darlinglab.org/mauve/mauve.html) and the genes contained in each block were marked on the plot.

### SSR analyses

Microsatellites were analyzed by MISA (Beier et al., 2017; https://webblast.ipk-gatersleben.de/misa) on the type 1 reference mitogenome of *T. esculentum*, looking for repeats of motifs one to six base pairs long. The minimum number of repetitions were set to 10, 6, 5, 5, 5, 5 for mono-, di-, tri-, tetra-, penta-, and hexanucleotide repeats, respectively.

### Phylogenetic tree construction

Two phylogenetic trees were constructed separately, one based on the mitogenome conserved gene sequences to explore the evolutionary relationship between *T. esculentum* and several other selected legumes, and the other tree was built on all differential loci found in the mitogenomes of *T. esculentum* to explore the inter-population and intra-population relationship among the 43 independent samples.

The 24 conserved mitochondrial genes, *atp1*, *atp4*, *atp6*, *atp8*, *atp9*, *nad3*, *nad4*, *nad4L*, *nad6*, *nad7*, *nad9*, *mttB*, *matR*, *cox1*, *cox3*, *cob*, *ccmFn*, *ccmFc*, *ccmC*, *ccmB*, *rps3*, *rps4*, *rps12*, and *rpl16* from the mitogenomes of *T. esculentum* (OK638188 and OK638189), *Arabidopsis thaliana* (NC_037304.1), *C. canadensis* (MN017226.1), *L. japonicus* (NC_016743.2), *M. sativa* (ON782580.1), *M. pinnata* (NC_016742.1), *G. max* (NC_020455.1), and *V. radiata* (NC_015121.1) were concatenated to make artificial chromosomes. A Maximum Likelihood (ML) phylogenetic tree using the Jukes-Cantor model was built in Mega 11 (Tamura et al., 2021; https://www.megasoftware.net/) after the chromosomes were aligned by Muscle v5 (Edgar, 2022; https://www.drive5.com/muscle/). The topology was validated by a Bayesian inference phylogenetic tree drawn by BEAST v1.10.4 (Suchard et al., 2017; https://beast.community/) and FigTree v1.4.4 (http://tree.bio.ed.ac.uk/software/figtree/).

Artificial chromosomes concatenated by 40 bp segments at each of the 254 differential loci found in the mitogenomes of *T. esculentum* were prepared for the 43 independent individuals and aligned by Muscle v5. A Maximum Likelihood (ML) phylogenetic tree using the Jukes-Cantor model was drawn on the 43 chromosomes. Frequencies from 1000 bootstrap replicates were labeled on the branches with 40% as cutoff. The topology was verified by a neighbor-joining tree in Mega 11.

### Genetic information exchange between organelles

The reference chloroplast genome sequence of *T. esculentum* (KX792933.1) was blasted to its type 1 reference mitogenome (OK638188 and OK638189) and visualized by the Advanced Circos function in TBtools (Krzywinski et al., 2009; Chen et al., 2020; https://github.com/CJ-Chen/TBtools). Primers were designed to verify the presence of the 9,798bp long homologous fragment in both the mitochondrial and chloroplast genomes. Four pairs of primers were designed, using NCBI Primer-BLAST, to amplify the products spanning the two ends of the 9,798 bp cpDNA insertion in the mitochondrial and chloroplast genomes, respectively. The two pairs of primers designed based on the mitogenome sequence included: MitoLL Forward (ACGCAGAAAAGAGGCCGAA) and MitoLR Reverse (CCTTCGTTTAAGAGAATGTTTTTGG), and the product size was 117 bp. MitoRL Forward (TCTTTGCTACAGCTGATAAAAATCG) and MitoRR Reverse (CCTATGTTCGTTTTCGCCCTG), and the product size was 120 bp. The two pairs of primers designed based on the plastome sequence included: ChlLL Forward (CGTAGTCGGTCTGGCCC) and MitoLR Reverse (CCTTCGTTTAAGAGAATGTTTTTGG), and the product size was 117 bp. MitoRL Forward (TCTTTGCTACAGCTGATAAAAATCG) and ChlRR Reverse (GCTTTTAATAATATGGCCGTGATCT), and the product size was 120 bp.

20 uL of mixtures were prepared by separately mixing DNA from two randomly selected *T. esculentum* samples and Promega GoTaq Green Master Mix. The amplifications started with denaturation at 95 °C for 5 min, followed by 32 cycles of 95 °C for 30 s, 55 °C for 30 s, and 72 ° C for 30 s, and a final 72 °C for 5 min.

## Results

### Genome structure and rearrangement

When the WGS Illumina reads from the 84 individuals were mapped to the two chromosomes of the reference mitogenome of *T. esculentum*, LS1 (OK638188) and LS2 (OK638189), two distinct mitogenomes were found. The mitogenomes of 45 individuals were similar to the reference mitogenome, with only a few substitutions and indels seen, termed type 1. However, the mitogenomes of the other 39 individuals were very different from the reference, with a large number of sequence and structural differences, termed type 2 (Table S1). This is consistent with the previously published study of *T. esculentum* chloroplast genomes, where these 84 individuals were found to contain two distinct germplasms (Li and Cullis, 2023). These two cytotypes actually differ not only in the chloroplast genome but also in the mitochondrial genome. The PacBio HiFi reads from the type 2 individual Sample 4 were assembled by Canu to generate three circular molecules M1 (OP795449), M2 (OP795450), and M4 (OP795447), and one linear chromosome M3 (OP795448), with a total length of 436,568 bp and a GC content of 44.8% (Table 1) (Figures S1-5). The four chromosomes consisted of 21 contigs assembled directly from Illumina reads of type 2 individuals, containing four double-copy regions and one triple-copy region (Figure 1 and Table 2). Among these multi-copy regions, homologous sequences of contigs H and I were also doubled in coverage in the type 1 mitogenome of *T. esculentum*, but the rest were present as single-copy sequences in the type 1 mitogenome (Li and Cullis, 2021).

**Figure 1.**
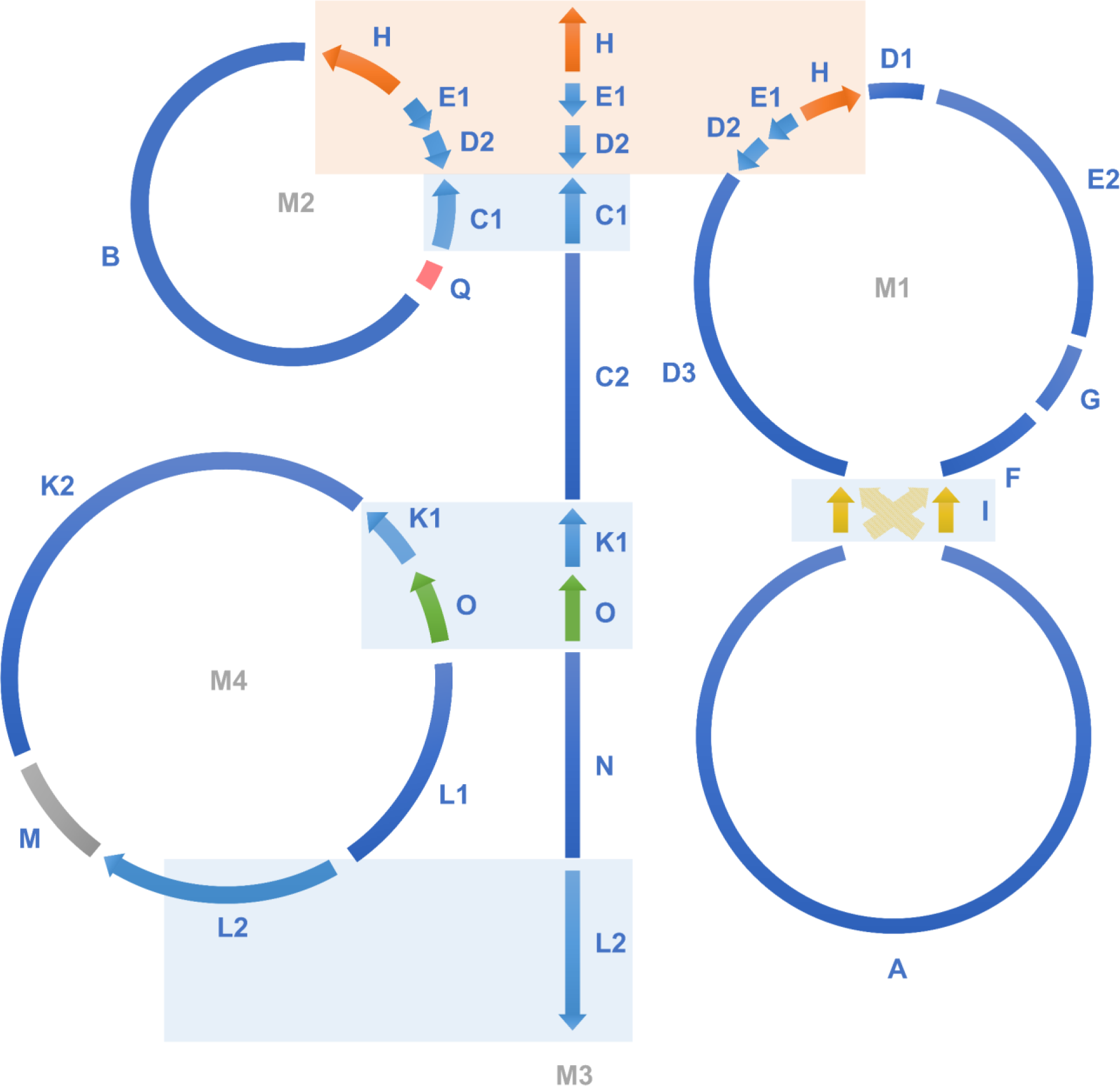
The assembly graph of the type 2 mitogenome of *T. esculentum*. The type 2 mitogenome consists of three circular molecules M1, M2, and M4, and one linear molecule M3, and they were built on 21 long scaffolds (Table 1). Blue blocks show double-copy regions that are identical between two chromosomes, and orange blocks indicate triple-copy regions owned by three chromosomes. Close sequencing coverage was found for single-copy regions of the four chromosomes. Recombination on a pair of long inverted repeats I can change the junction of the upper and lower halves of M1 to that indicated by the yellow dashed arrows. The two structures before and after recombination have been confirmed by PacBio long reads, and their frequencies were close in the same individual (Figure S6-9). A long chloroplast insertion was found at the position of the gray segment M, about 9,798 bp, and its length varied slightly among different individuals. This chloroplast insertion and long repeats H, I, and O are also present in the type 1 mitogenome of *T. esculentum*. The type 1 mitogenome also has two rings, very similar to the M1 and M4 here, but other molecules have undergone dramatic changes.

**Table 1.** Chromosome base composition of the type 2 T. esculentum mitogenome.

| Molecule | A (%) | C (%) | G (%) | T (%) | G~C (%) | Length (bp) |
| --- | --- | --- | --- | --- | --- | --- |
| M1 | 27.90 | 22.18 | 22.22 | 27.70 | 44.40 | 169,406 |
| M2 | 27.60 | 21.63 | 23.26 | 27.52 | 44.90 | 56,355 |
| M3 | 27.46 | 22.25 | 22.73 | 27.55 | 45.00 | 97,120 |
| M4 | 27.70 | 22.58 | 22.44 | 27.29 | 45.00 | 113,687 |

**Table 2.** Length of the primary scaffolds constituting the type 2 T. esculentum mitogenome.

| Unit number | Length (bp) | Unit number | Length (bp) |
| --- | --- | --- | --- |
| A | 82,874 | K1 | 4,052 |
| B | 39,273 | K2 | 52,730 |
| C1 | 30,017 | L1 | 20,069 |
| C2 | 5,272 | L2 | 22,301 |
| D1 | 5,866 | N | 27,581 |
| D2 | 2,722 | O | 4,881 |
| D3 | 26,671 | M | 9,654 |
| H | 5,212 | Q | 2,108 |
| E1 | 1,768 |  |  |
| E2 | 25,384 |  |  |
| F | 8,590 |  |  |
| G | 5,798 |  |  |
| I | 2,265 |  |  |

Both ends of the chromosome M3 were found to be long repeats that were homologous to parts of other chromosomes (Figure 2). In addition, in a very long range of 15 to 20 kb, the sequencing depth gradually decreased towards both ends, and many reads were found to stop in this range. This furthers confirms that this is a linear chromosome that exists in different lengths in cells because of the lack of telomere protection. Linear chromosomes have been found to stably exist in eukaryotic cells even in the absence of telomeres, through strand-invasion between terminal sequences and their homologous internal sequences to form t-loops to protect the chromosomes from degradation (de Lange, 2015). Because the repeats at both ends of M3 are very long, the PacBio reads we obtained cannot span them to verify whether this linear molecule recombines with other chromosomes. Long range PCR amplification that can amplify sequences above 20 kb can be considered here to answer this question.

**Figure 2.**
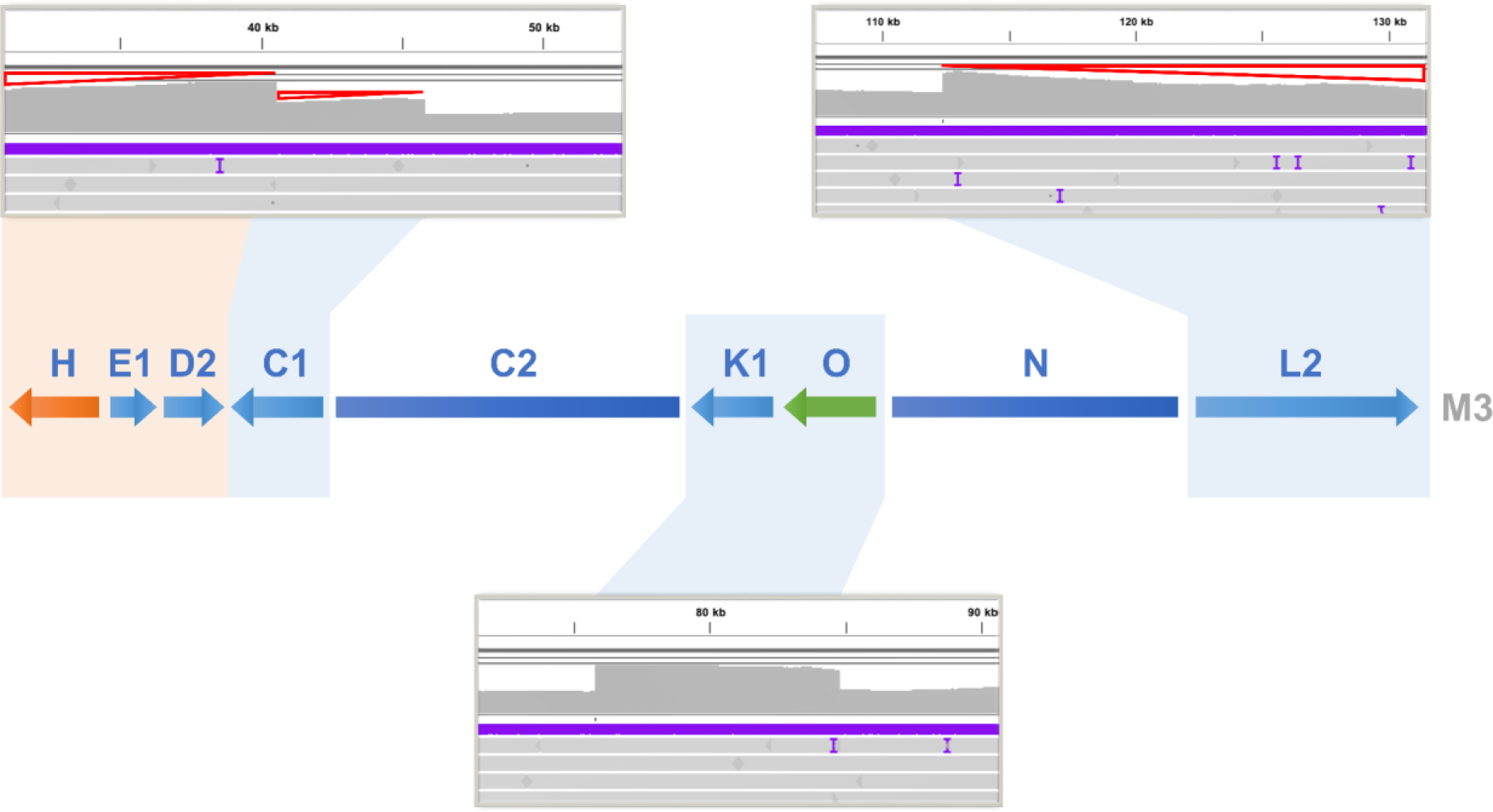
Changes in sequencing coverage on the type 2 mitogenome chromosome M3 of *T. esculentum*. PacBio HiFi reads from the type 2 individual Sample 4 were aligned to the multicopy regions of the chromosome M3 using pbmm2. Chromosome M3 contains some long repeats that are identical to parts of other chromosomes, thus increasing the sequencing coverage of these regions in the alignment (Figures S10-12). At the positions of scaffolds C1, K1-O, and L2, with blue shading, the read depth was found to be doubled. At the location of scaffold H shaded in orange, the sequence depth was increased to 3-fold. However, a progressive decrease in coverage indicated by red triangles was also seen at both ends of M3, as some linear M3 chromosomes had degenerated at both ends without telomere protection.

Numerous differences were found between the type 1 and type 2 mitogenome structures (Figure 3). One is a rare recombination on a pair of *trnfM* genes on the large circular molecule M1, which inverts the 12,846 bp sequence in between. The sequence before inversion also appeared in type 2 plants, with a frequency less than 2%. Furthermore, the three small rings of the type 1 mitogenome exist in different forms in the type 2 mitogenome, and four gaps have been found on them. New type 2 exclusive fragments were discovered, including a 2,108 bp segment, which connected originally distantly located contigs C1 and B. In addition, the recombination on a pair of 35 bp direct repeats joined contigs C1 and D2 to form a new circular molecule M2. C2 was found to connect with K1 and then further extended to N and L2 to form a linear chromosome, but the mechanism behind this is unclear. It can be seen that the deletion and insertion of the entire DNA segment, alongside repeat mediated recombination, can lead to dramatic changes in the mitogenome structure.

**Figure 3.**
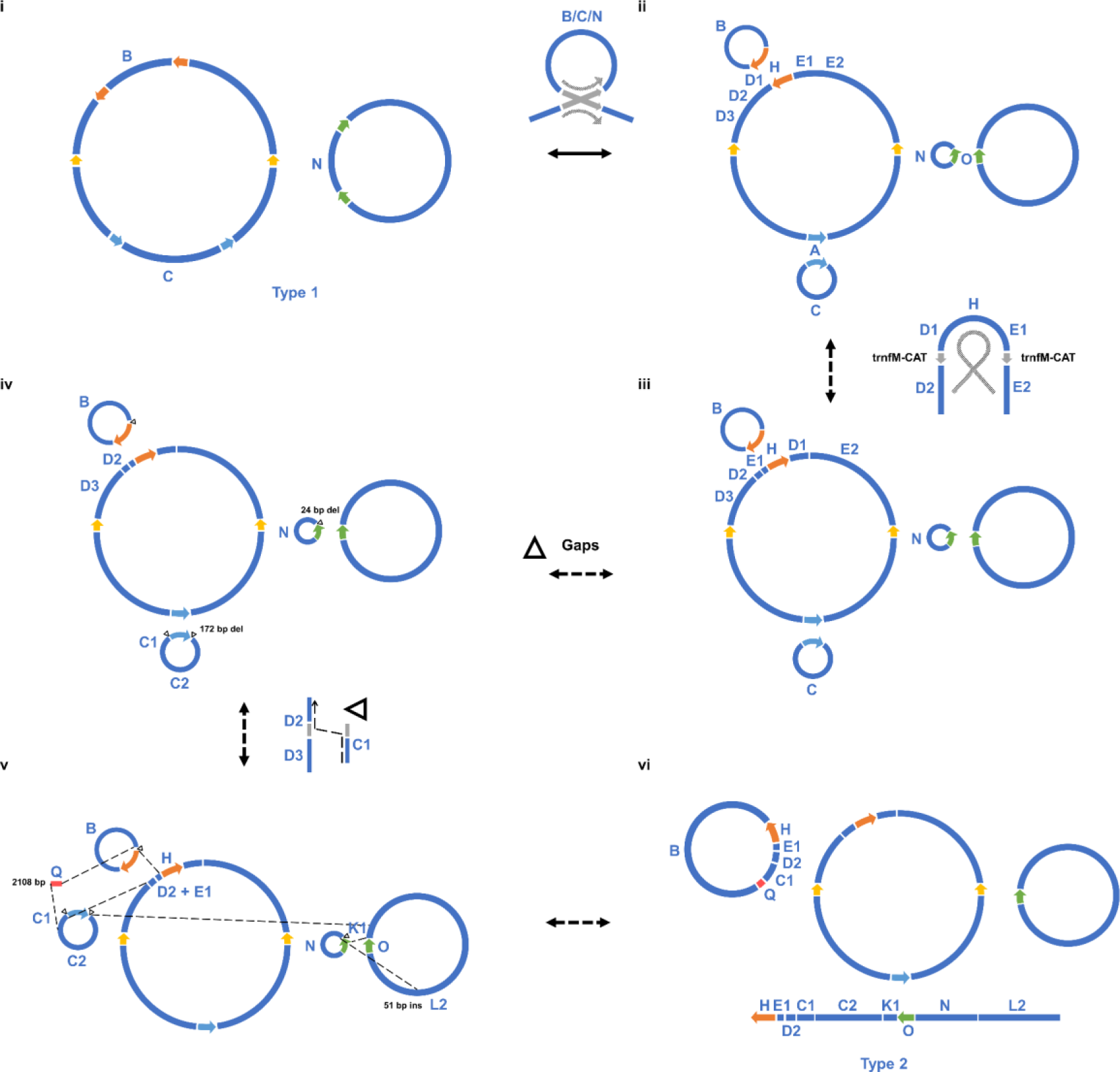
Step-by-step analysis of the structural differences between the two types of *T. esculentum* mitogenomes. i. The two autonomous circular chromosomes of type 1 *T. esculentum* mitogenome, LS1 (OK638188) (left) and LS2 (OK638189) (right). Colored arrows indicate the four pairs of long repeats (>1 kb). ii. Recombination on the direct repeats split the two large rings into five small circular molecules. Both conformations before and after recombination have been confirmed by PacBio reads to exist in type 1 individuals. iii. A rare recombination on a pair of *trnfM* genes at the junctions of D1 and D2, and E1 and E2, inverted the sequence D1-H-E1 in between. iv. Gaps with or without sequence deletions resulted in the three small circular chromosomes of the type 1 mitogenome present as different forms in type 2 individuals (Figures S13-18). v. New DNA fragments, including a 2,108 bp contig Q unique to type 2 individuals, joined originally remotely located sequences to form new structures. vi. The final type 2 mitogenome of *T. esculentum* with three circular and one linear chromosomes.

The two types of *T. esculentum* mitogenomes were compared by NUCmer alignment and then visualized by a synteny diagram, showing a high degree of similarity (Figure 4). The basic blocks making up the two mitogenomes are highly similar, except for the 2,108 bp type 2 unique fragment and some other short insertions and deletions, but the order of these blocks has been changed by recombination. When the WGS Illumina reads from the 84 individuals were mapped to the region where the 2,108 bp type 2 unique fragment resides, 39 samples were found to contain this fragment while 45 plants did not (Figures S19-28). Furthermore, in type 1 mitogenome, LS1 and LS2 are two autonomous circular chromosomes that do not recombine into one master circle, but in type 2 mitogenome, a linear chromosome M3 was found to contain homologous sequences from both LS1 and LS2, suggesting that these two molecules may have been related in evolution.

**Figure 4.**
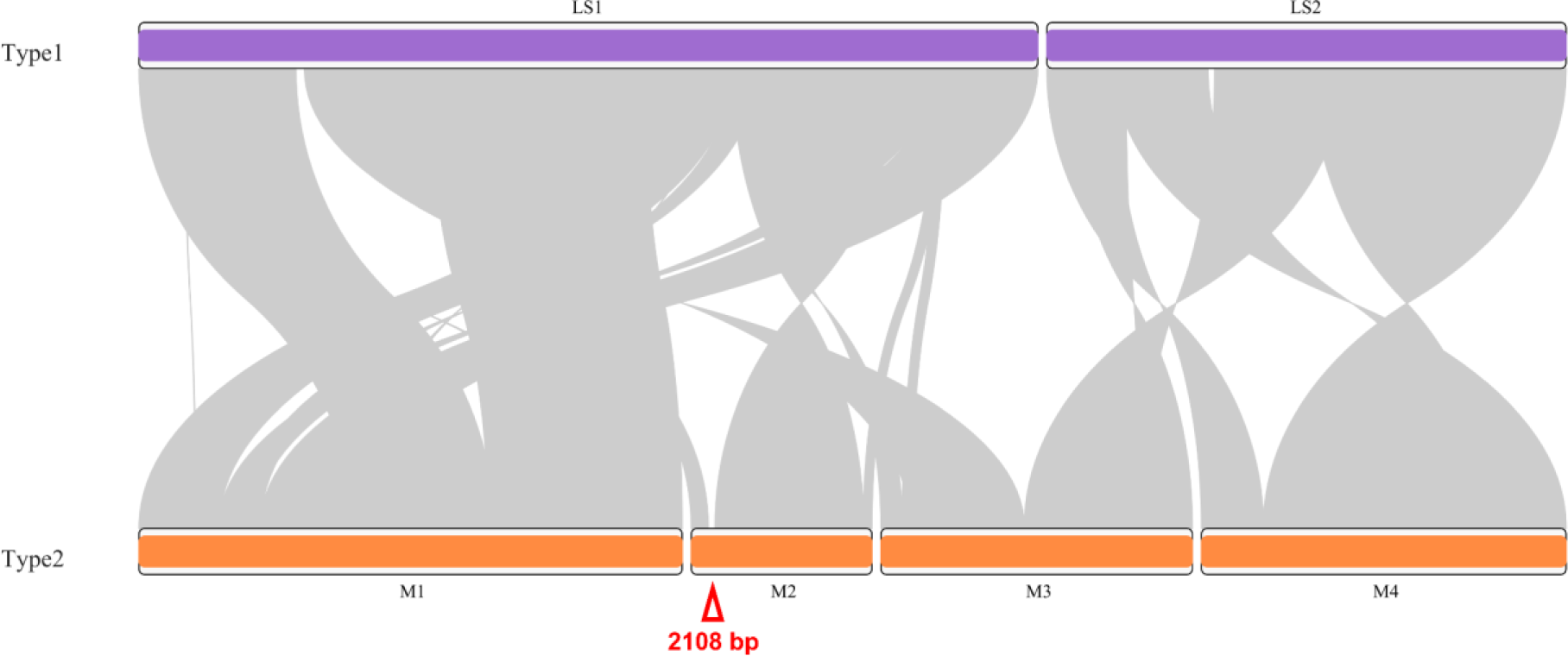
Synteny visualization of the two types of mitogenomes of *T. esculentum* by the R package RIdeogram after NUCmer alignment. The red triangle indicates the 2,108 bp type 2 mitogenome exclusive fragment of *T. esculentum*.

Blast results showed that this 2,108 bp type 2 mitogenome exclusive fragment was highly similar to the mitochondrial sequences of Fabaceae species *Lupinus albus* and *Indigofera tinctoria*, suggesting that this fragment was possibly derived in the evolution from *Lupinus* or *Indigofera* (Figure 5A). Two pairs of primers were designed and found to effectively identify the 2,108 bp fragment. As shown in Figure 5B and 5C, sample B and C, out of the six randomly selected samples, contained both the 990 bp left end band and the 289 bp right end band after amplification, indicating that only these two of the six had type 2 mitogenomes.

**Figure 5.**
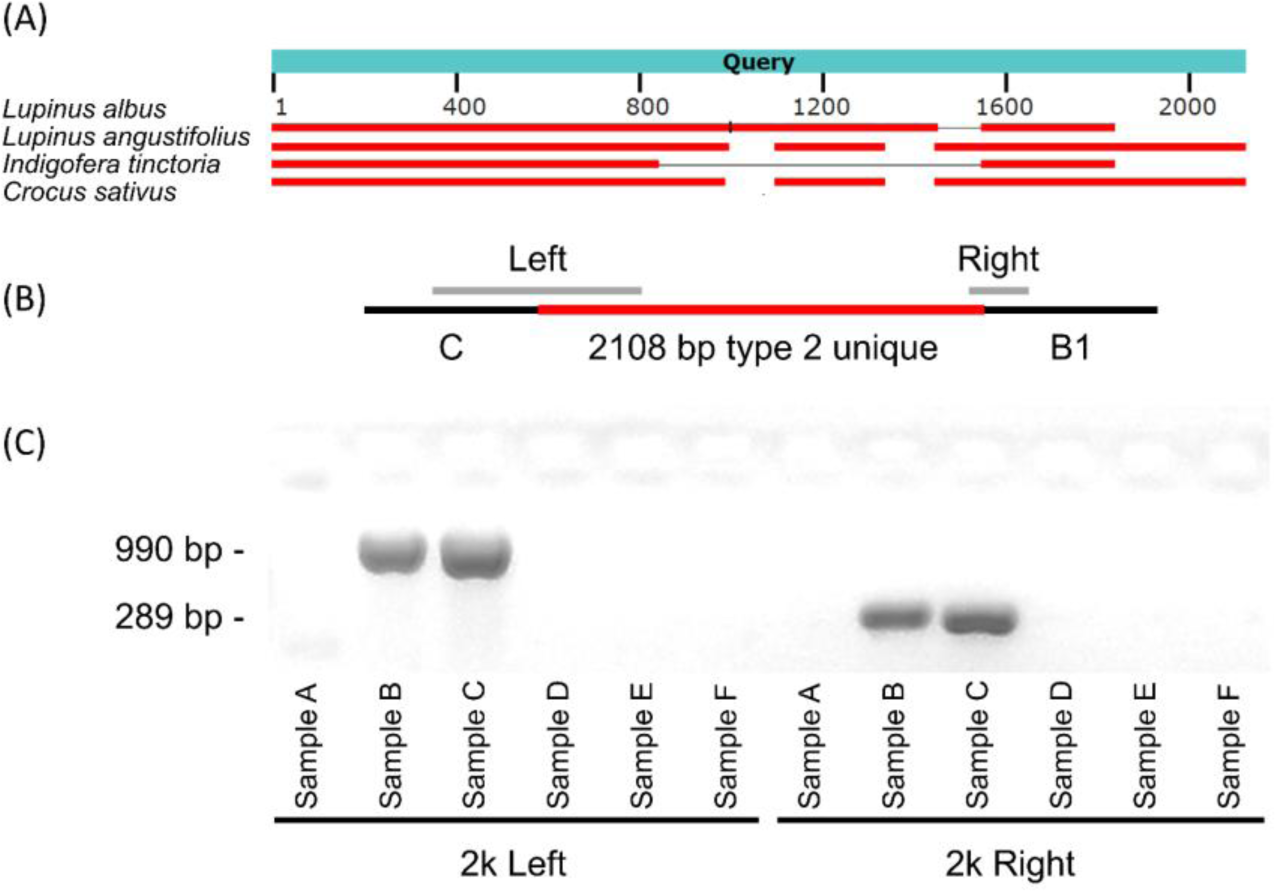
Homology analysis of the 2,108 bp fragment unique to type 2 *T. esculentum* mitogenome and design of primers for its PCR identification. (A) The 2,108 bp type 2 mitogenome exclusive sequence was blasted as a query in the NCBI database. Red horizontal bars indicate where database sequences are aligned, and separately aligned regions from the same database subject are connected by thin gray lines. (B) Two pairs of primers were designed to amplify products across both ends of the 2,108 bp fragment. The estimated size of the left end product is 990 bp, and the right end product is 289 bp. (C) Gel image of PCR amplification of DNA from six randomly selected samples with the two pairs of primers designed separately. The PCR products were electrophoresed on a 1.5% agarose gel at 80V for 1 hour.

### Gene annotation

Both type 1 and type 2 mitogenomes of *T. esculentum* were found to contain 35 unique protein-coding genes, 3 unique rRNA genes, and 16 different tRNA genes (Figure 6; Table 3) (Li and Cullis, 2021). The type 2 mitogenomes have two copies of *nad9* and *atp8*. The gene *nad9* is located on contig K1, a long repeat possessed by both chromosomes M3 and M4 in type 2 mitogenomes. Whereas, there is only one copy of K1 in type 1 mitogenomes. The gene *atp8* is located on a pair of long repeats J, so its copy number is doubled as is the case in both types of mitogenomes. The copy number of exon 3 and 4 of gene *nad5* is also doubled in type 2 mitogenomes but not in type 1, and it is not known whether this affects its expression level. In addition, there are two copies of *rrn5* and *rrnS* in type 2 mitogenomes but only one copy in type 1. A total of 26 tRNA genes were found in type 2 mitogenomes, including four copies of *trnfM-CAT*, three copies of *trnM-CAT*, three copies of *trnC-GCA*, two copies of *trnP* and *trnQ*, and 12 single-copy tRNA genes.

**Figure 6.**
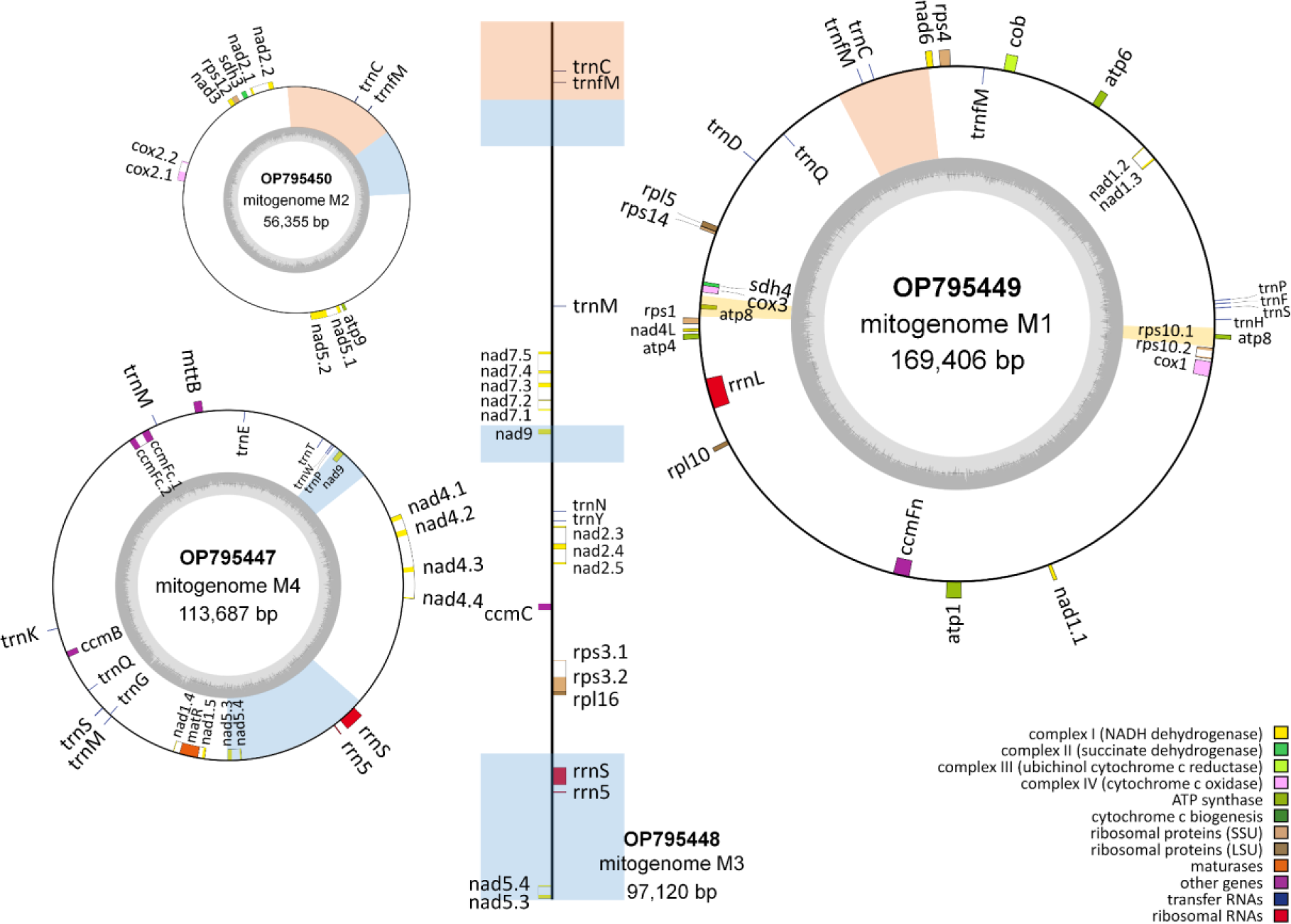
The map of type 2 *T. esculentum* mitogenome gene arrangement drawn by OGDRAW. The annotation was performed by MITOFY and BLAST on the mtDNA of individual Index1 from UP Farm and deposited in GenBank under accession numbers OP795447-OP795450. All type 2 plants were found to have a similar gene arrangement. The dark gray pattern in the inner circle indicates GC content. Genes are colored according to their function. The decimal part after the gene name indicates the order of the exons. Genes inside the circle are transcribed clockwise, while those outside the circle are transcribed counterclockwise. Blue blocks represent two-copy regions that are identical between two chromosomes, and orange blocks show three-copy regions that are the same across three chromosomes.

**Table 3.** Gene annotation of the type 2 mitogenome of *T. esculentum*.

| Category | Gene Name |  |  |  |
| --- | --- | --- | --- | --- |
|  | M1 | M2 | M3 | M4 |
| Complex I (NADH dehydrogenase) | *nad1#*, *nad4L*,<br>*nad6* | *nad2#*, *nad3*,<br>*nad5#* | *nad2#*, *nad5#*,<br>*nad7#*, *nad9* | *nad1#*, *nad4#*,<br>*nad5#*, *nad9* |
| Complex II (succinate dehydrogenase) | *sdh4* | *sdh3* |  |  |
| Complex III (ubiquinol cytochrome-c reductase) | *cob* |  |  |  |
| Complex IV (cytochrome-c oxidase) | *cox1, cox3* | *cox2#* |  |  |
| Complex V (ATP synthase) | *atp1, atp4, atp6, atp8\*(2)* | *atp9* |  |  |
| Cytochrome c biogenesis | *ccmFn* |  | *ccmC* | *ccmB, ccmFc#* |
| Large subunit ribosomal proteins | *rpl5, rpl10* |  | *rpl16* |  |
| Small subunit ribosomal proteins | *rps1, rps4, rps10#, rps14* | *rps12* | *rps3#* |  |
| Maturases |  |  |  | *matR* |
| Transport membrane protein |  |  |  | *mttB* |
| Ribosomal RNAs | *rrnL* |  | *rrn5, rrnS* | *rrn5, rrnS* |
| Transfer RNAs | *trnD-GTC, trnC-GCA, trnQ-TTG, trnH-GTG, trnfM-CAT\*(2), trnF-GAA, trnP-TGG, trnS-GCT* | *trnC-GCA, trnfM-CAT* | *trnN-GTT, trnC-GCA, trnfM-CAT, trnM-CAT, trnY-GTA* | *trnQ-TTG, trnE-TTC, trnG-GCC, trnK-TTT, trnM-CAT\*(2), trnP-TGG, trnS-TGA, trnT-TGT#, trnW-CCA* |

Genes with introns are marked with #. Genes with multiple copy numbers on the same chromosome are labeled with \*, and the numbers in parentheses indicate the corresponding copy numbers.

The atypical start codon ACG was used by three genes *nad1*, *nad4L*, and *rps10*, and ATT was used by gene *mttB*. This is consistent with the research on the mitogenome of common beans from which these four genes were all reported to use an alternative initiation codon ACG (Bi et al., 2020). C-to-U editing was found to be widely used in mitochondrial and chloroplast genes in land plants (Takenaka et al., 2013). ATT is also usually used as an alternative start codon in the mitogenome. For example, *mttB* in *Salix purpurea* was reported to use an ATT start codon as well (Wei et al., 2016).

### Mitogenome divergence

The mitogenome of *T. esculentum* was highly divergent from those of the six selected legume species, which covered 23% to 48% of the marama mitogenome, ranging in length from 91.9-191.8 kb (Figure 7). Of these species, *C. canadensis* was most closely related to marama, while *M. sativa* was the least similar to marama. The mitogenomes of *G. max*, *L. japonicus*, and *V. radiata* all contain homologous sequences covering 25% of the marama mitogenome, equal to 99.89 kb in length. *Bauhinia variegata* is closer to marama than *C. canadensis* in the phylogenetic tree, but its mitogenome sequence is not available in NCBI GenBank (Wunderlin, 2010).

**Figure 7.**
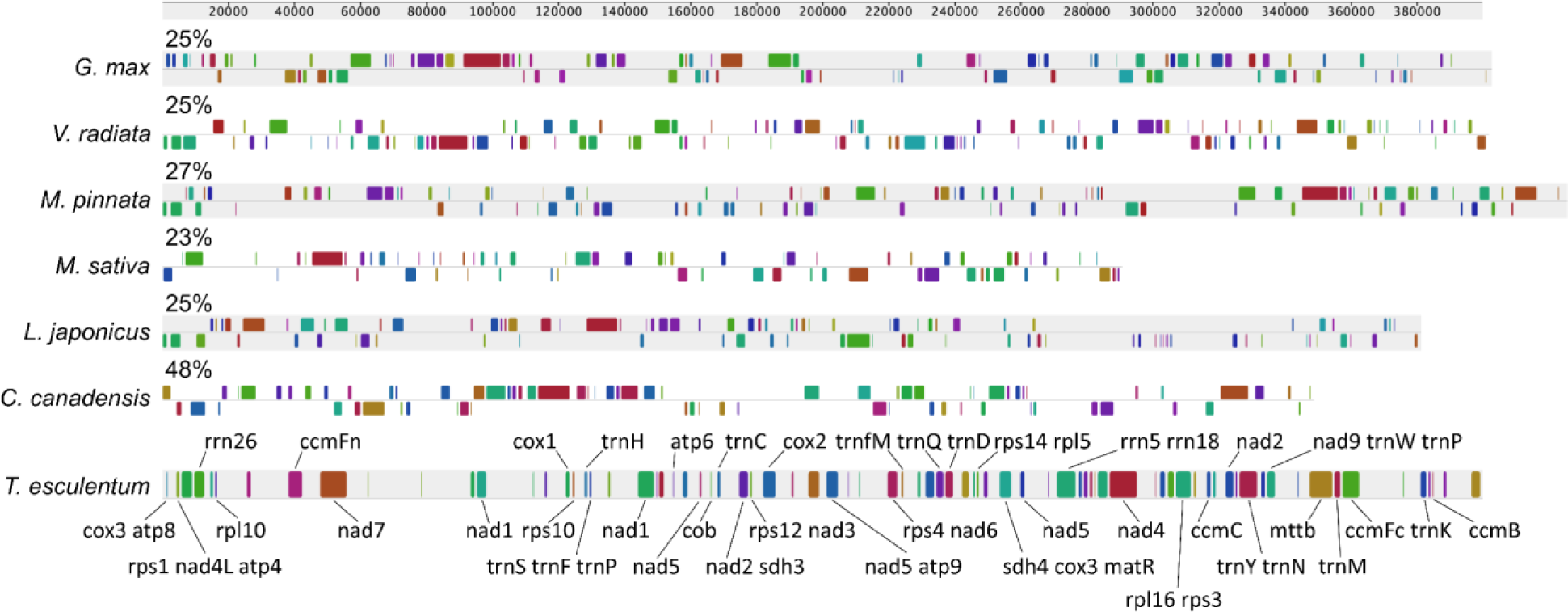
Synteny block diagram of the Mauve alignment between the mitogenomes of *T. esculentum* and six other Fabaceae species, *Cercis canadensis* (MN017226.1), *Lotus japonicus* (NC_016743.2), *Medicago sativa* (ON782580.1), *Millettia pinnata* (NC_016742.1), *Glycine max* (NC_020455.1), and *Vigna radiata* (NC_015121.1). Similarities (percentage of marama mitogenome covered) are labeled above the bars. Genes contained in the synteny blocks are marked by gene symbols.

The loss of mitochondrial protein-coding genes and the functional transfer of these genes from the organelle genome to the nuclear genome are common in the evolution of angiosperms, but some plants tend to retain a more complete set of mitochondrial genes (Palmer et al., 2000). The mitogenomes of Cercidoideae species *T. esculentum* and *C. canadensis* contain functional protein-coding genes *sdh3*, *sdh4*, and *rpl10*, which have been lost in many other legumes (Table 4). Rare gene losses were also seen in the mitogenomes of legumes, such as *rpl5* in *G. max*, *cox2* in *V. radiata* and *rps1* in *L. japonicus*, but these genes all remain intact and functional in *T. esculentum* and *C. canadensis* (Alverson et al., 2011; Kazakoff et al., 2012; Chang et al., 2013).

**Table 4.** List of mitochondrial protein-coding genes lost during the evolution of some Fabaceae species.

| Gene | *T. esculentum* | *C. canadensis* | *L. japonicus* | *M. sativa* | *M. pinnata* | *V. radiata* | *G. max* |
| --- | --- | --- | --- | --- | --- | --- | --- |
| *sdh3* | + | + | - | - | + | - | - |
| *sdh4* | + | + | - | - | - | - | - |
| *cox2* | + | + | + | + | + | - | + |
| *rpl2* | - | - | - | - | - | - | - |
| *rpl5* | + | + | + | + | + | + | - |
| *rpl10* | + | + | - | - | - | - | - |
| *rps1* | + | + | - | + | + | + | + |
| *rps2* | - | - | - | - | - | - | - |
| *rps7* | - | - | - | - | - | - | - |
| *rps8* | - | - | - | - | - | - | - |
| *rps11* | - | - | - | - | - | - | - |
| *rps13* | - | - | - | - | - | - | - |
| *rps19* | - | - | - | - | - | - | - |

In a pairwise comparison of the mitogenomes of seven legume species, the synteny plot revealed numerous rearrangements and a high degree of divergence among the mitogenomes of even closely related species (Figure 8). The mitogenomes of *C. canadensis* and *T. esculentum* contain many distinct regions but also some long homologous segments.

**Figure 8.**
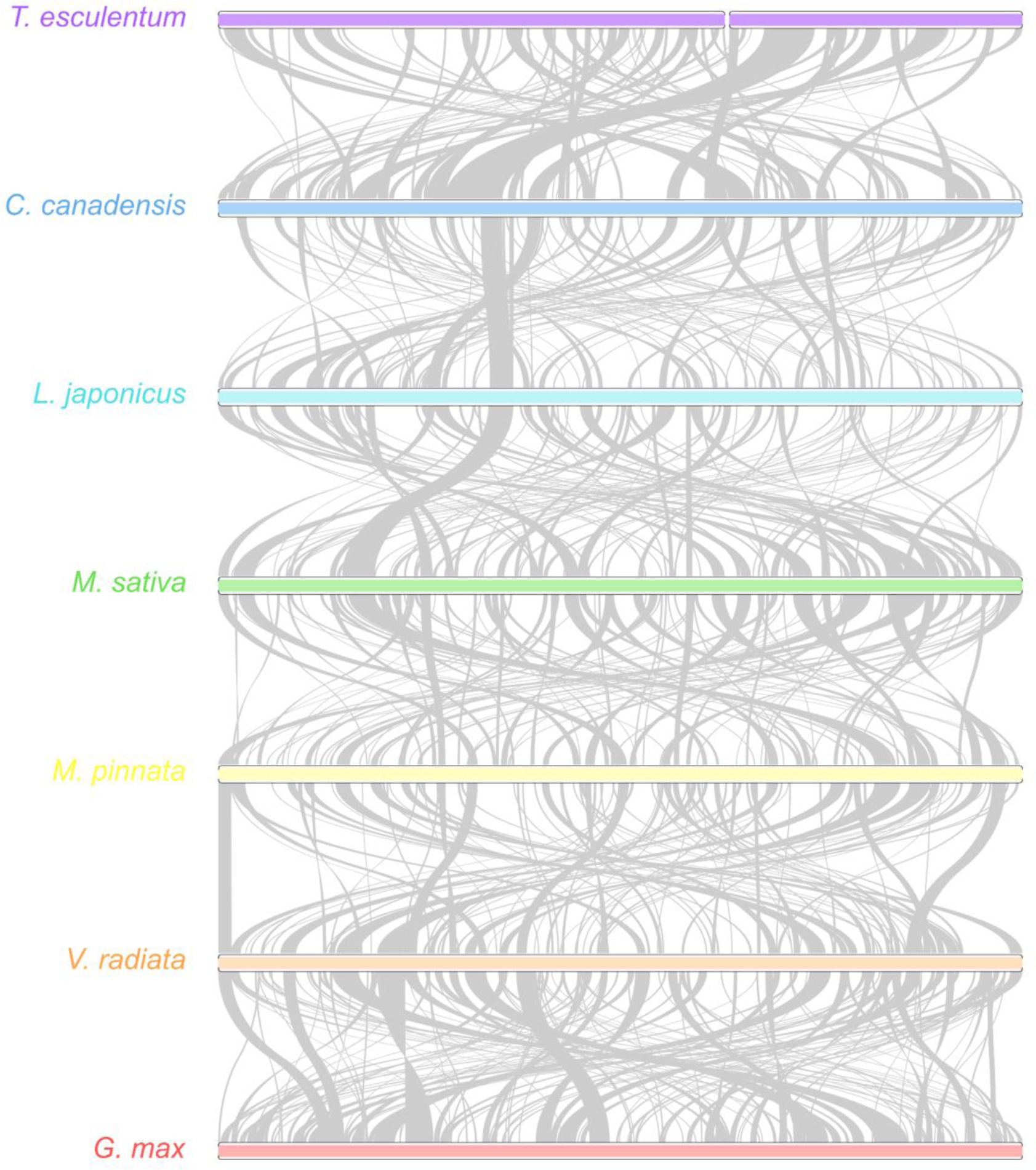
Synteny maps of the mitochondrial genomes of seven legume species. The colored bars represent the mitochondrial chromosomes of *C. canadensis* (MN017226.1), *L. japonicus* (NC_016743.2), *M. sativa* (ON782580.1), *M. pinnata* (NC_016742.1), *V. radiata* (NC_015121.1), *G. max* (NC_020455.1), and *T. esculentum*, which contains two chromosomes, LS1 (OK638188) and LS2 (OK638189). The gray ribbons indicate homologous sequences between the two neighboring species. The species were ordered according to the phylogenetic tree in Figure 9. Promer was used to detect syntenic regions between highly divergent genomes, which were then visualized by RIdeogram package in R.

The phylogenetic tree shown in Figure 9 was built on the 24 conserved mitochondrial protein-coding genes *atp1*, *atp4*, *atp6*, *atp8*, *atp9*, *nad3*, *nad4*, *nad4L*, *nad6*, *nad7*, *nad9*, *mttB*, *matR*, *cox1*, *cox3*, *cob*, *ccmFn*, *ccmFc*, *ccmC*, *ccmB*, *rps3*, *rps4*, *rps12*, and *rpl16*, which are present in all these eight species. This tree is consistent with previously published phylogenetic trees constructed on chloroplast protein-coding genes (Kim and Cullis, 2017; Wang et al., 2018). As another species of Cercidoideae, *C. canadensis* was expected to be the closest relative of these plants to *T. esculentum*. Among Faboideae species, *M. sativa* and *L. japonicus* are closely related, and *V. radiata*, *G. max*, and *M. pinnata* belong another clade.

**Figure 9.**
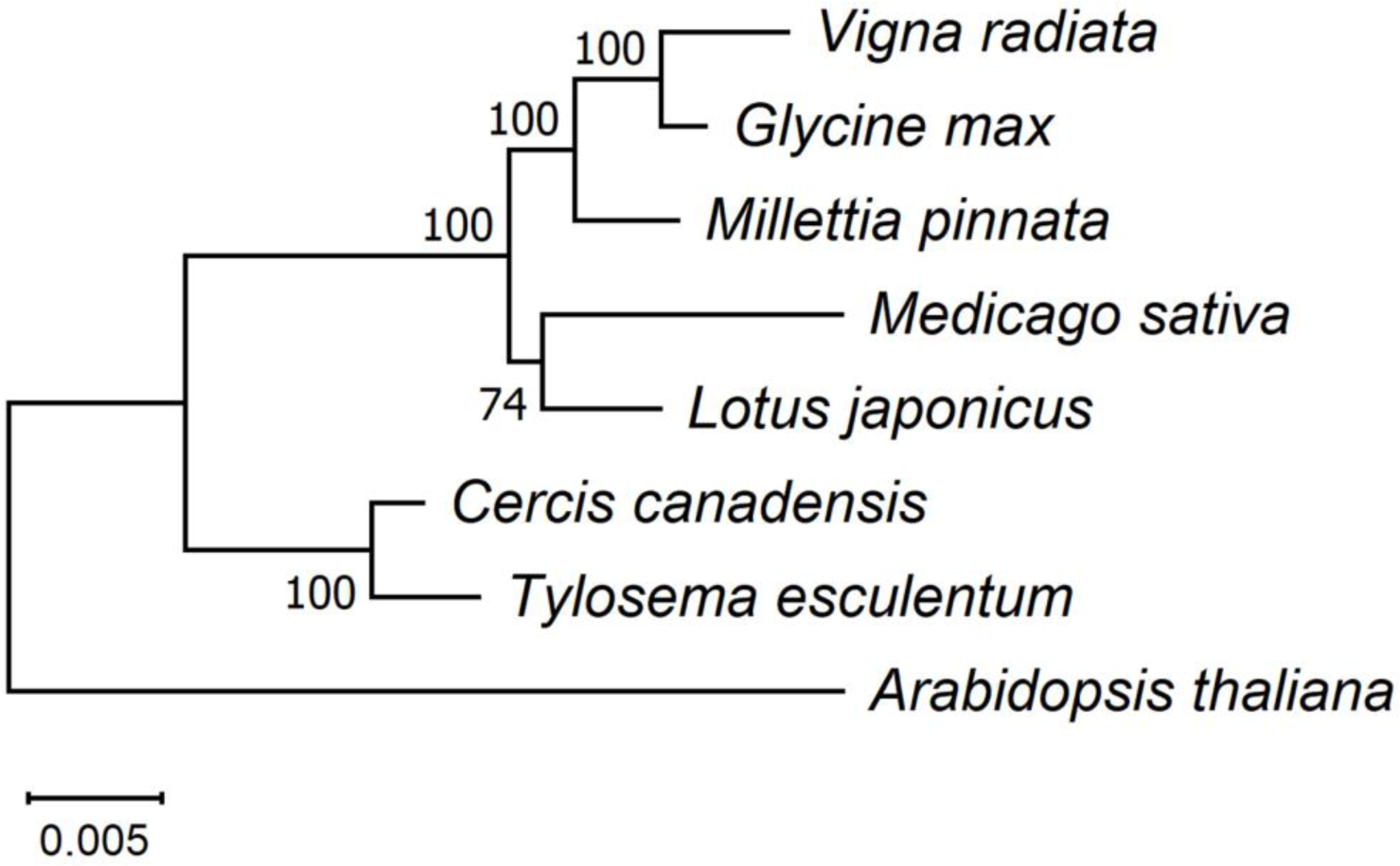
Maximum Likelihood (ML) phylogenetic tree with the Jukes-Cantor model based on artificial chromosomes concatenated by 24 conserved mitochondrial genes, *atp1*, *atp4*, *atp6*, *atp8*, *atp9*, *nad3*, *nad4*, *nad4L*, *nad6*, *nad7*, *nad9*, *mttB*, *matR*, *cox1*, *cox3*, *cob*, *ccmFn*, *ccmFc*, *ccmC*, *ccmB*, *rps3*, *rps4*, *rps12*, and *rpl16* from *Arabidopsis thaliana* (NC_037304.1), *Cercis canadensis* (MN017226.1), *Lotus japonicus* (NC_016743.2), *Medicago sativa* (ON782580.1), *Millettia pinnata* (NC_016742.1), *Glycine max* (NC_020455.1), and *Vigna radiata* (NC_015121.1) in NCBI. The tree was drawn in Mega 11 after sequence alignment with Muscle v5. Percentage probabilities based on 1000 bootstrap replications are labeled on the branches. The topology was validated by the Bayesian inference phylogenetic tree drawn by BEAST (Figure S29).

### Nucleotide polymorphism

17 haplotypes were found in these 47 plants, which could be clearly divided into two groups: namely the type 2 plants from the Namibian farm and the UP farm and the remaining type 1 plants (Figure 10). The mitogenomes of type 2 plants are relative conserved, which may be caused by sampling errors, and a larger sample size is needed to verify. The only differences between type 2 plants were four deletions at four closely located loci on the chromosome LS2. However, the type 1 plants can be divided into many groups according to the variations. Some geographical patterns can be seen in the distribution of variation. For example, in the four plants from Osire, there were three substitutions on the chromosome LS1, A>C at 146,140 bp, C>A at 173,053 bp, and C>A at 217,508 bp, and a deletion at 128,192 bp on the chromosome LS2. These are variations unique to Osire plants. Similar geographic-specific variation can be seen in plants elsewhere. However, our data may still have sampling issues. For example, only a single sample was collected in some regions including Otjiwarongo and Okamatapati. In addition, although distant wild individuals in each geographic region were intentionally selected, there is no guarantee that they are not related. The findings here still need to be validated by sequencing more samples and studied alongside the phenotypic performance of the plants to determine whether any of these variations are the result of plants evolving to better adapt to different environments.

**Figure 10.**
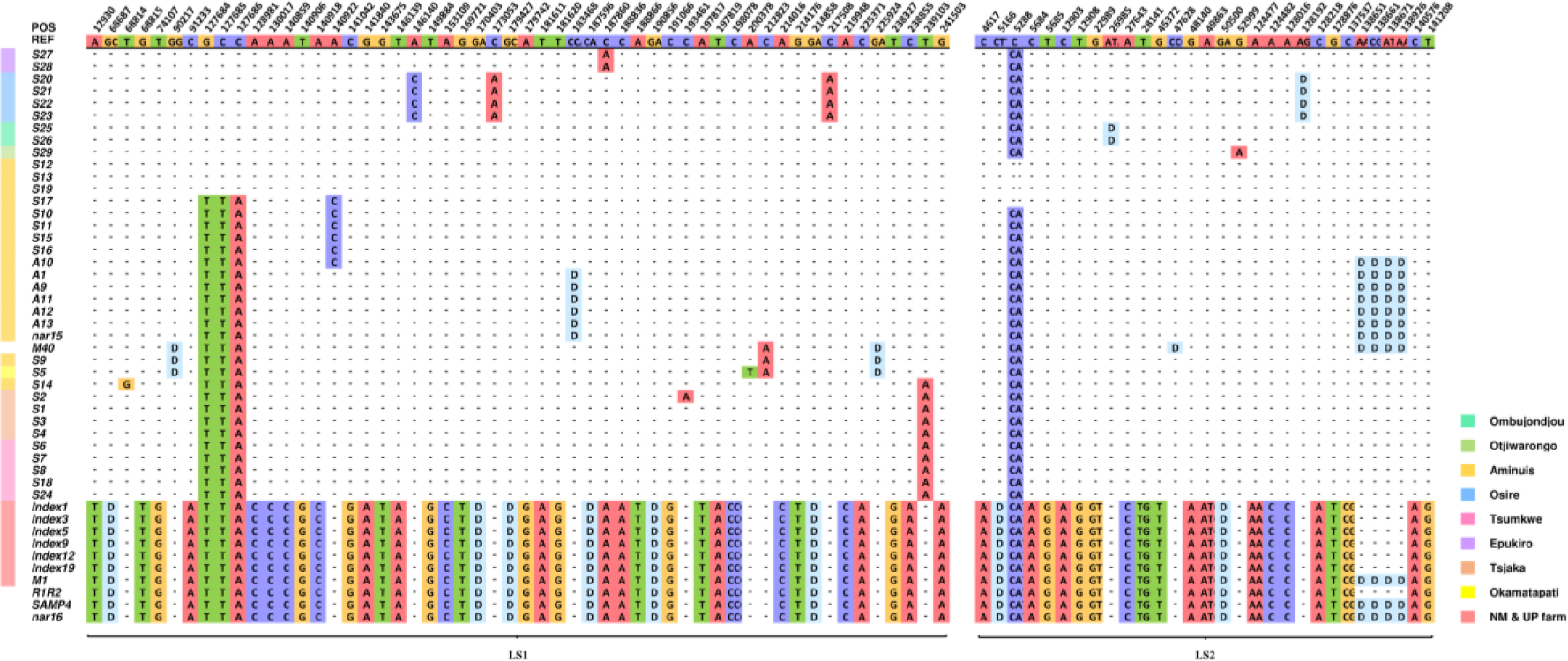
Nucleotide matrices showing the distribution of mitochondrial genome variations in the 43 independent individuals and 4 additional samples of unknown origin. The first row indicates the alleles of the type 1 reference mitogenome of *T. esculentum* (LS1:OK638188 and LS2: OK638189). From the second row onwards, only bases different from the reference are shown, and bases identical to the reference are replaced by dashes. The two types of mitogenomes also differ from each other at another 170 loci, not shown here to save space (no within-type differences were found at those 170 loci). The full variation distribution is shown in Figure S30 and S31. All insertions are represented by the first two bases and deletions by the letter “D”. The color bar to the left of the plant ID shows the source of the sample and is left blank for unknown sources.

Type 2 refers to 7 Index plants excluding Index8 from the Pretoria Farm, 29 M descendent plants originally from the Namibia Farm excluding M40, and two individuals R1R2 and nar16 of unknown origin. Type 1 represents all remaining plants, including A plants, S plants, Index8, and M40. Numbers in parentheses indicate counts of exclusive variants of this type. LS1 and LS2 are type 1 marama reference mitochondrial chromosomes in GenBank with accession numbers OK638188 and OK638189.

A total of 254 differential loci were found in the mitogenomes of the 84 *T. esculentum* individuals, including 143 SNPs, 52 insertions, and 59 deletions (Table 5). Type 1 and type 2 mitogenomes differed at 230 loci, including 129 substitutions, 50 deletions, and 51 insertions. The mitogenomes of type 2 plants differed at only 4 loci, whereas that of type 1 plants differed at 24 loci.

**Table 5.** Total number of variations found when mapping the WGS reads of all 84 individuals to the type 1 *T. esculentum* reference mitochondrial genomes OK638188 and OK638189.

| Variation Type | Type 2 vs. Ref. | Type 1 vs. Ref. | Total |
| --- | --- | --- | --- |
| **LS1** |  |  |  |
| Deletion | 29 (29) | 3 (3) | 32 |
| Insertion | 34 (34) | 0 | 34 |
| SNP | 79 (75) | 13 (9) | 88 |
| **LS2** |  |  |  |
| Deletion | 25 (21) | 7 (3) | 27 |
| Insertion | 18 (17) | 1 (0) | 18 |
| SNP | 54 (54) | 1 (1) | 55 |
|  | 239 (230) | 24 (16) | 254 |

The mitochondrial gene sequence of *T. esculentum* is very conserved. A total of 11 variations were found in the mitochondrial gene sequence, and only 1 of them was in the coding sequence, which was a 2368A>G substitution and resulted in a N303D change in the gene *matR* (Table 6). Furthermore, 10 of the 11 variations were found on one subgenomic ring LS2 of the reference mitogenome of *T. esculentum*. Whether the chromosome LS1 is more conserved than the chromosome LS2 is unknown. Although the intergenic spacer of LS1 contained more variations than LS2, the gene sequence on LS1 appeared to be more conserved, and cpDNA insertions were also found rarely in LS1, but abundantly in chromosome LS2.

**Table 6.**
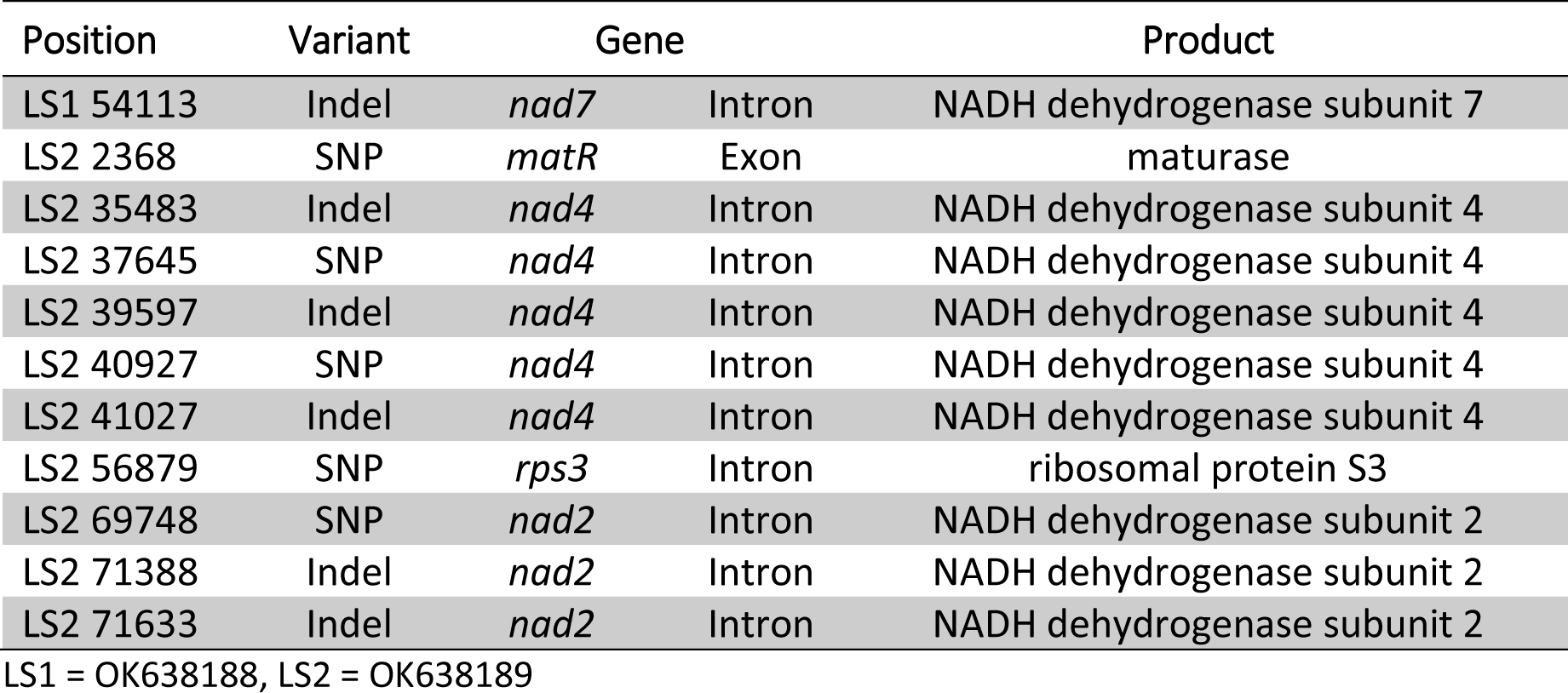
Variations found in T. esculentum mitochondrial gene sequences of the 84 individuals.

In the phylogenetic tree constructed on the differential loci of the mitogenomes of *T. esculentum*, the two germplasms fell into two clusters as expected (Figure 11). Furthermore, the type 1 plants were then divided into groups. Tsumkwe and Tsjaka were two distant sampling sites, but the plants from these two locations were clustered in one clade, which was also seen in the phylogenic tree built on the complete chloroplast genome. One wondered whether there were factors other than geographical distance that determined the grouping of these plants. On the other hand, plants from Epukiro, Osire, Ombujondjou, and Otjiwarongo belonged to one clade, and these sites were geographically close together and were all arid areas with soil moisture anomalies (SMA) typically below −.04 m^3^/m^3^ (NASA, n.d.). In contrast, plants from another clade seemed to grow in less arid regions. Notable differences between the mitogenomes of plants from the two clades included three consecutive substitutions from GCC to TTA at positions 127,684 to 127,686 on chromosome LS1 (OK638188), although the function is unknown. Of course, the sample size of this study is still relatively small, and more plants need to be collected in areas with different soil moisture for verification.

**Figure 11.**
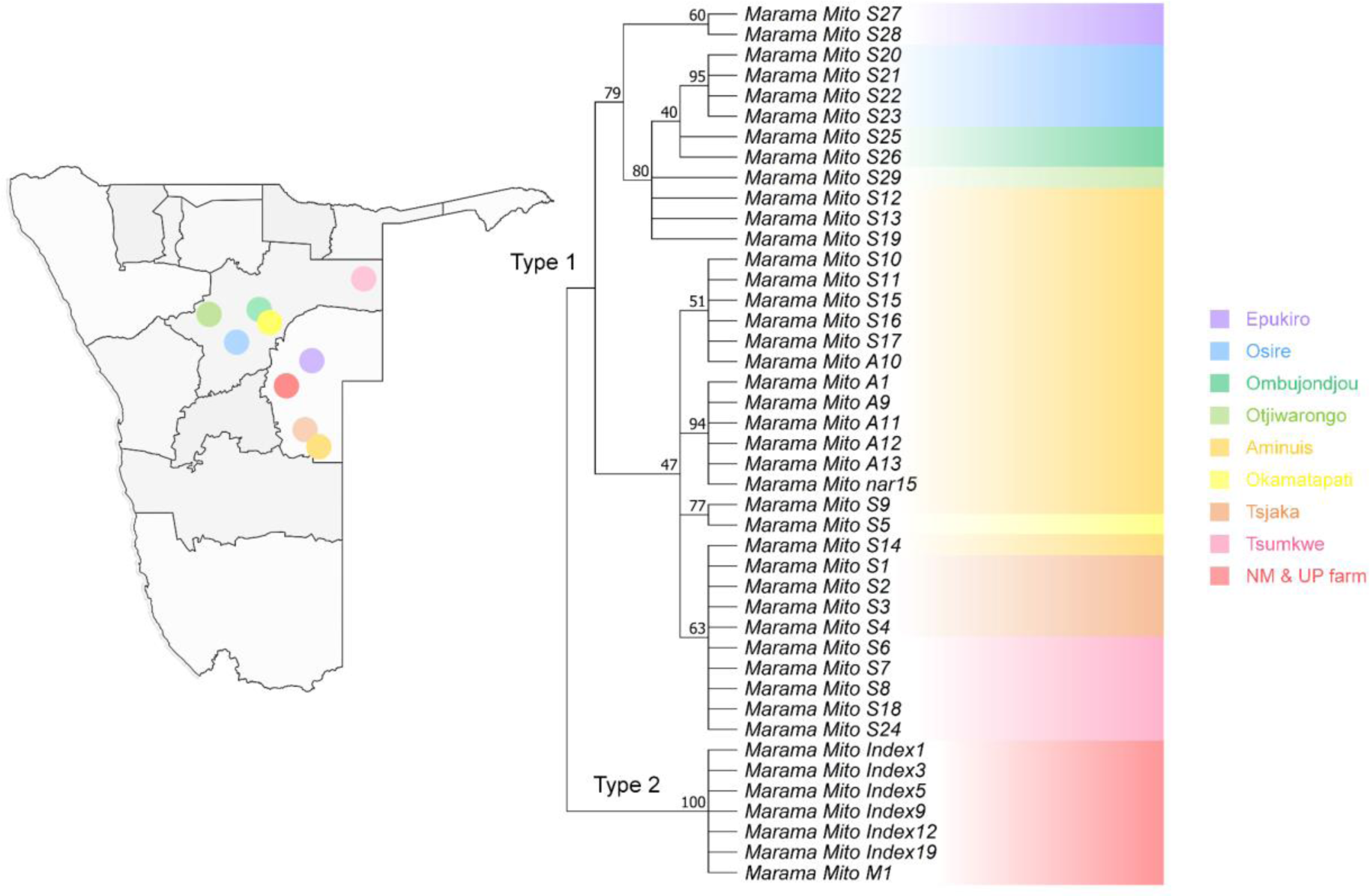
Maximum Likelihood (ML) phylogenetic tree with the Jukes-Cantor model built on artificial chromosomes concatenated by 40 bp fragments at each of the 254 differential loci in the mitogenomes of *T. esculentum* according to the mitogenome sequences of the 43 independent individuals. Frequencies from 1000 bootstrap replicates were labeled on the branches with 40% as cutoff. The topology was verified by the neighbor-joining method in Mega 11. Individuals with the same background color came from the same geographic location, and the sampling points were marked on the map of Namibia.

### SSRs and heteroplasmy analyses

A total of 48 SSR motifs were found by MISA in the reference mitogenome of *T. esculentum*, LS1 (OK638188) and LS2 (OK638189), of which 38 were simple mononucleotide microsatellites, accounting for 79.2% of all discovered SSR motifs (Figure 12). Among them, 37 are A/T mononucleotide repeats, and only one is a G/C repeat. There are 8 dinucleotide repeats and 2 trinucleotide repeats. No simple sequence repeats with core motifs of four nucleotides or longer were found. There are three microsatellites in the coding sequence of the gene, including two 10 bp A/T repeats, one located at the boundary of the coding sequence of the gene *sdh3*, the other located in the exon 4 of the gene *nad1*, and a 12 bp AT repeat siting in the coding sequence of *mttB*.

**Figure 12.**
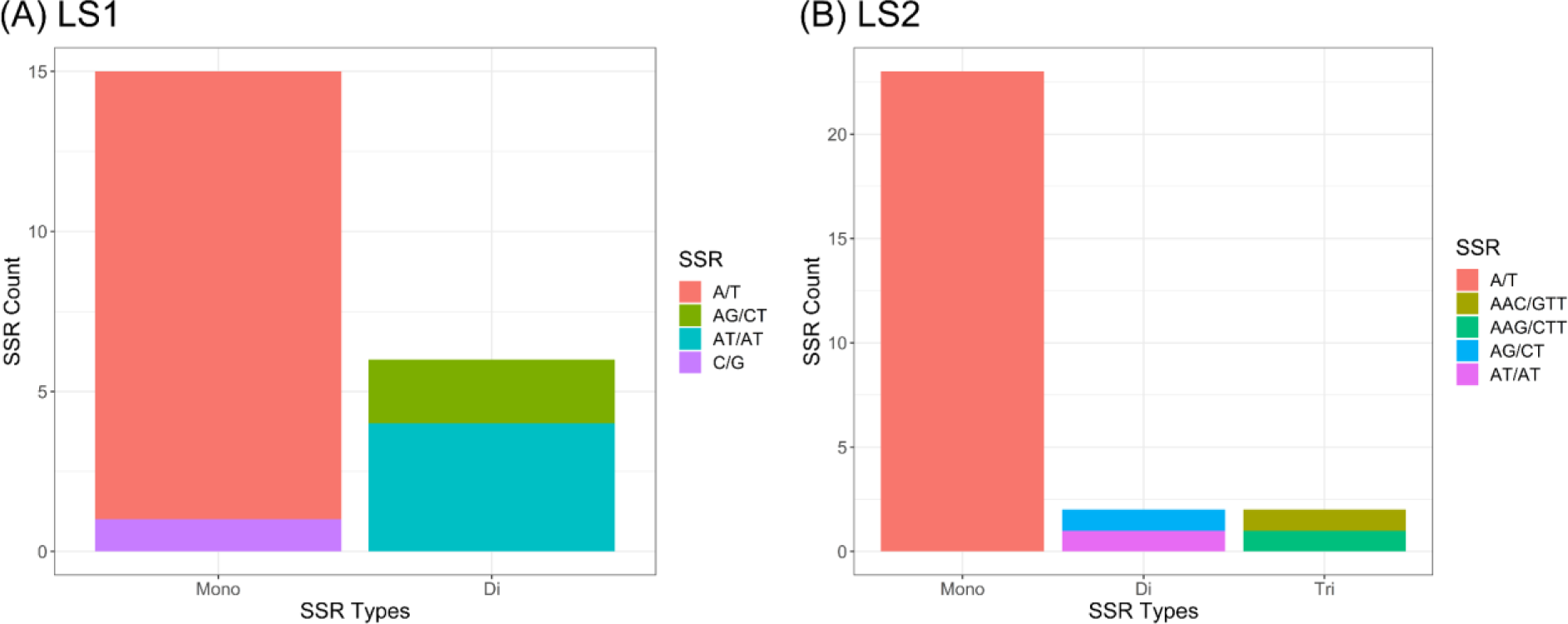
Distribution of SSR motifs of different repeat types in the type 1 reference mitogenome of *T. esculentum* analyzed by MISA. (A) Chromosome LS1 (OK638188). (B). Chromosome LS2 (OK638189).

An individual collected from Aminuis, A11 had approximately 96% type 1 alleles and 4% type 2 alleles at all differential loci between the two types of marama mitogenomes (Figure 13). It has been confirmed that these minor alleles are not from homologous sequences in the nuclear genome or the chloroplast genome, suggesting the presence of two different mitogenomes in the same individual. It was previously reported that heteroplasmy also exists in the chloroplast genome of A11, with a minor genome frequency reaching 11% (Jin and Cullis, 2023). This means that in the chloroplasts and mitochondria of A11, both major and minor genomes exist, and the frequency of the minor genome is higher than that of the other studied plants. This is most likely to be caused by an accidentally occurred paternal leakage. The proportions of mitochondrial and chloroplast minor genomes differed in A11, indicating that this may be true heterogeneity rather than due to accidental mixing of samples. However, being the only sample with high overall organelle genome heterogeneity is not convincing, and more plants from Aminuis need to be sequenced and studied.

**Figure 13.**
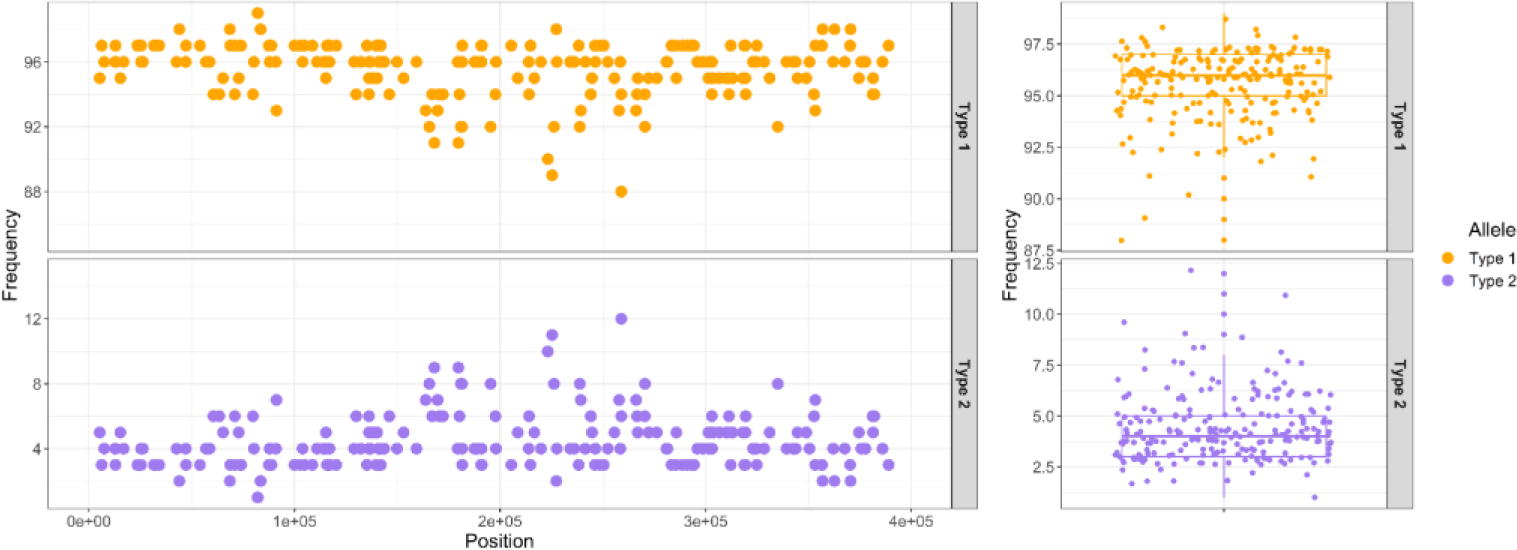
Allele frequency plot of all differential loci between the two types of mitogenomes of *T. esculentum* in Aminuis individual A11. The x-axis of the left panel indicates the position on the chromosome concatenated by the reference marama mitogenomes LS1 (OK638188) and LS2 (OK638189).

In the mitogenomes of other individuals, heteroplasmy was found to be less common than in their chloroplast genomes, and even low proportions of base substitutions below 2% were relatively rare, and even if present, many were found from mitochondrial homologous segments in the nuclear genome. However, at a few differential loci, including the three consecutive substitutions at positions 127,684 to 127,686 and another substitution at 140,922 on chromosome LS1 (OK638188), obvious heteroplasmy could be seen in multiple individuals, with the proportion of minor alleles even up to 35%. These loci may have played important roles in the evolution of these individuals under environmental selection.

### Sequence transfer between chloroplast and mitochondrial genomes

It was interesting to find that the chloroplast DNA insertions were concentrated in a subgenomic ring of the mitogenome (Figure 14). A low collinearity between the two genomes was seen, and the inserted cpDNA fragments seemed to be completely rearranged. These contained a 9,798 bp fragment, in which a large number of variations were observed. Primers were designed to amplify across the two ends of this fragment in both plastome and mitogenome to verify its presence (Figure 15). 20 variations were found in this long homologous DNA fragment, of which 13 mutations occurred in the mitogenome and 7 occurred in the chloroplast genome (Table 7). These do not include the differences between the two organelle genomes at another 72 loci in this segment (this included 22 deletions, 6 insertions, and 44 SNPs.), which are the same for both germplasms, making it difficult to tell where the mutation occurred. 10 variations were found in the gene sequence on this segment, of which only 2 synonymous substitutions occurred in the chloroplast genome, and the remaining 8 were in the mitogenome, which had a great impact on transcription, including the introduction of early stop codons (Table S3).

**Figure 14.**
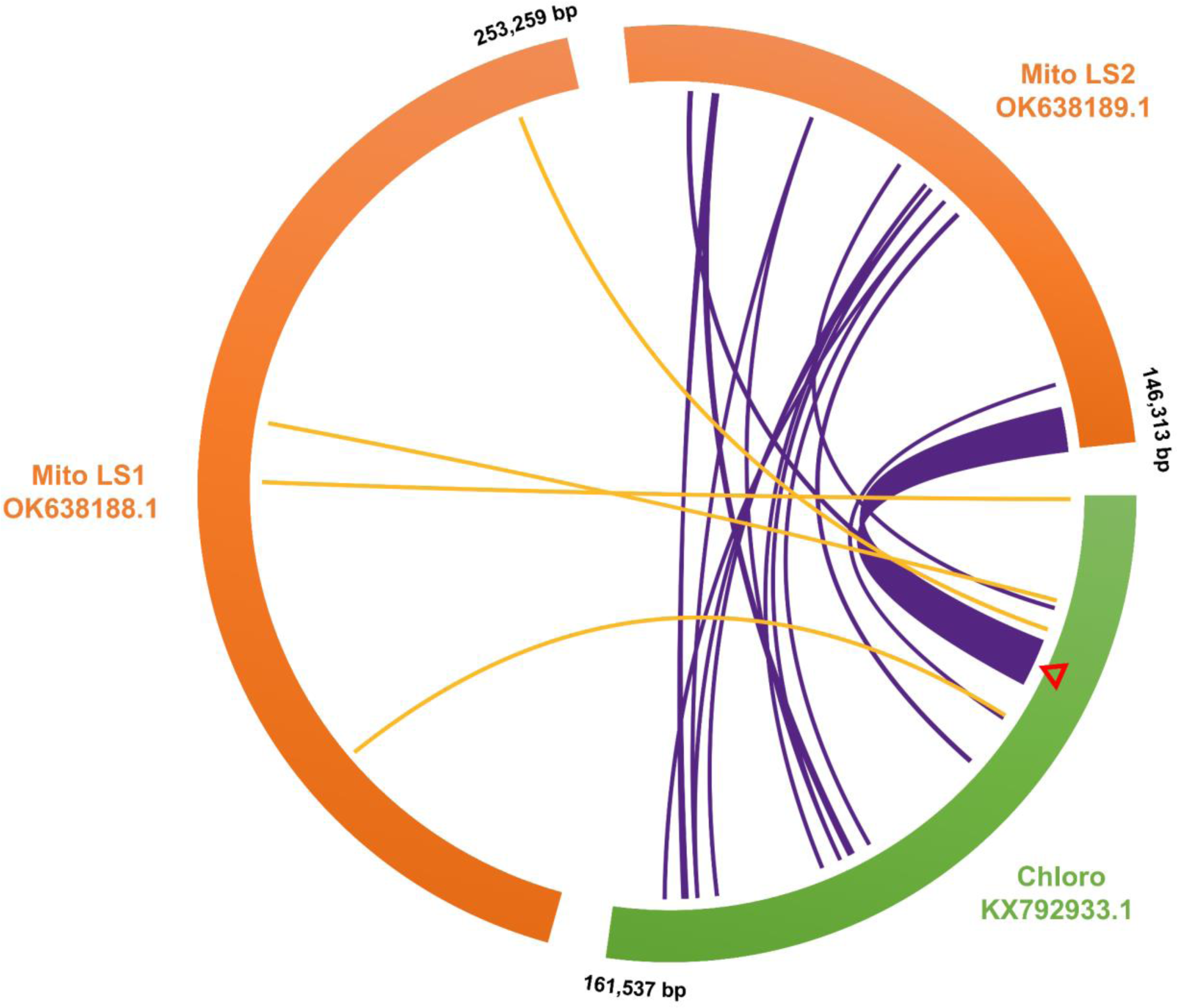
Map of chloroplast DNA insertions in the mitochondrial genome of *T. esculentum* drawn by TBtools Advanced Circos. The reference mitogenome chromosomes LS1 (OK638188) and LS2 (OK638189) were aligned with the chloroplast genome (KX792933) of *T. esculentum* using BLAST (Figure S32; Table S2). The curves in the middle connect the homologous sequences of the plastome and mitogenome. The cpDNA insertions in the two mitochondrial chromosomes are colored orange and purple, respectively. cpDNA insertions were concentrated on the mitochondrial chromosome LS2, with only four short fragments on LS1. All the three chromosomes are circular but are shown here as linear, with numbers next to them indicating their length and orientation. The red triangle marks the position of the 9,798 bp long homologous fragment.

**Figure 15.**
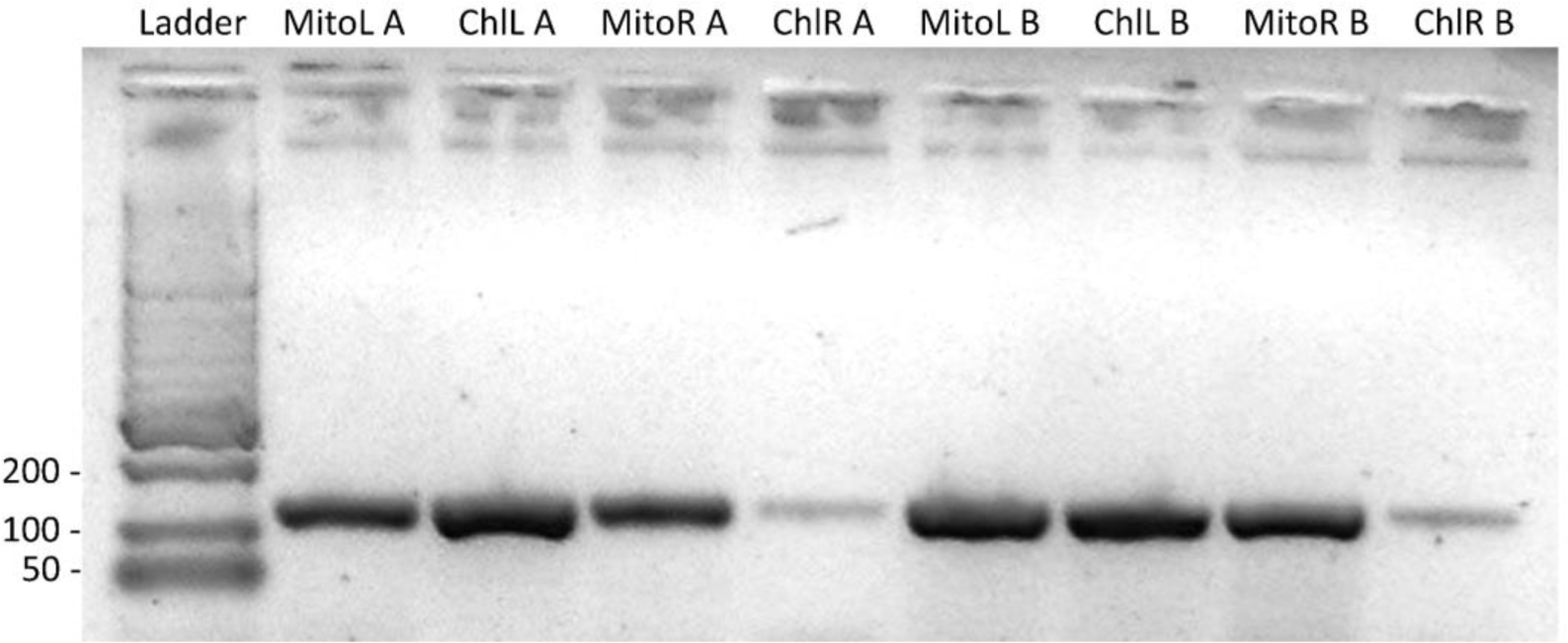
Amplification across the two ends of the 9,798 bp homologous fragment of the mitochondrial and chloroplast genomes in two random samples A and B. Lane1, HyperLadder II; Lanes 2-5, DNA from sample A; Lanes 6-9, DNA from sample B. The products in lanes 2 and 6 were amplified with primers MitoLL and MitoLR, in lane 3 and 7 were amplified with primers ChlLL and MitoLR, in lanes 4 and 8 were amplified with primers MitoRL and MitoRR, and in lanes 5 and 9 were amplified with primers MitoRL and ChlRR. The gel was run on a 1.5% agarose gel for 40 min at 100V. The annealing temperature of 55 °C was a bit high for the primer ChlRR, resulting in low yields and faint bands in lanes 5 and 9. Decreasing the annealing temperature by 1 °C and repeating the PCR resulted in normal amplification.

**Table 7.** Variations found in the 9,798 bp homologous segment within the organelle genomes of the 84 *T. esculentum* individuals.

| Position | Variation | Type 1 Chloro | Type 1 Mito | Type 2 Chloro | Type 2 Mito | Localization |
| --- | --- | --- | --- | --- | --- | --- |
| 35570* | SNP | Ref | **Alt** | Ref | Ref | Mito |
| 36347* | DEL | Ref | **Alt** | Ref | Ref | Mito |
| 36942 | SNP | Ref | Ref | **Alt** | Ref | Chloro |
| 37257 | SNP | **Ref** | Alt | Alt | Alt | Chloro |
| 37813 | SNP | **Alt** (some plants) | Ref | Ref | Ref | Chloro |
| 38753* | DEL | Ref | Ref | Ref | **Alt** | Mito |
| 38975 | SNP | Ref | Ref | **Alt** | Ref | Chloro |
| 39429* | SNP | Ref | **Alt** | Ref | Ref | Mito |
| 40061* | SNP | Ref | **Alt** | Ref | Ref | Mito |
| 40544* | SNP | Ref | **Alt** | Ref | Ref | Mito |
| 41243* | SNP | **Ref** | Alt | Alt | Alt | Chloro |
| 41716* | SNP | Ref | Ref | Ref | **Alt** | Mito |
| 41718* | SNP | Ref | Ref | Ref | **Alt** | Mito |
| 42587* | SNP | Ref | Ref | **Alt** | Ref | Chloro |
| 43559 | SNP | Ref | Ref | Ref | **Alt** | Mito |
| 44059 | INS | **Ref** | Alt | Alt | Alt | Chloro |
| 44235 | DEL | Ref | **Alt** | Ref | Ref | Mito |
| 44302 | SNP | Ref | **Alt** | Ref | Ref | Mito |
| 44552 | SNP | Ref | Ref | Ref | **Alt** | Mito |
| 44762 | DEL | Ref | Ref | Ref | **Alt** | Mito |

Refs refer to alleles identical to the type 1 chloroplast reference genome sequence. Alt refers to an allele that differs from the type 1 chloroplast reference genome sequence. Only two alleles, Ref or Alt, were seen at each locus. Comparing alleles in the same row, the locations where the mutations occurred can be estimated and written in the last column. This table does not include differences between plastome and mitogenome at another 72 loci on this fragment (no inter-type differences were seen at these loci). Positions in the gene sequence are marked with an asterisk\*.

The mitogenome of *T. esculentum* was 399,572 bp (type 1) in length, and a total of 254 variations were found in the mitogenomes of the 84 individuals. The length of the chloroplast genome was 161,537 bp (type 1), and a total of 147 variations were found in the plastomes of these plants. The value of variation per nucleotide of the chloroplast genome is even higher than that of the mitogenome, so it is speculated that these chloroplast genes *psbC*, *rps14*, *psaB*, and *psaA* are protected by some mechanism in the chloroplast genome, but after being transferred to the mitogenome, the protection mechanism disappears and these genes begin to accumulate mutations that render them nonfunctional and pseudogenes.

## Conclusion

The comparative analysis of the organelle genomes of 84 *T. esculentum* individuals revealed two germplasms with distinct mitochondrial and chloroplast genomes. The type 1 mitogenome contains two autonomous rings, or five smaller subgenomic circular molecules. These two equimolar structures are thought to be interchangeable through recombination on three pairs of long direct repeats (Li and Cullis, 2021). The type 2 mitogenome contains three circular molecules and one linear chromosome. It also has a unique fragment of 2,108 bp in length, likely derived from the mitogenome of *Lupinus*. Primers were designed to amplify on this fragment for germplasm typing. Both ends of the linear chromosome are repetitive sequences, also present in other molecules, on which recombination may occur to protect the linear chromosome from degradation.

The structural variation resulted in increased copy number of the genes *atp8*, *nad5* (exon3 and exon4), *nad9*, *rrnS*, *rrn5*, *trnC*, and *trnfM* in the type 2 mitogenome, but it is unknown whether this is reflected in the gene expression level. The genes *nad1*, *nad4L*, *rps10* and *mttB* were found to use alternative start codons ACG and ATT, similar to the mitogenome of common bean (Bi et al., 2020). A total of 254 differential loci were found in the mitogenomes of the 84 *T. esculentum* individuals. Type 1 and type 2 mitogenomes differ from each other at 230 of these loci. Only one of these 254 variations was found in the mitochondrial gene coding sequence, which altered the amino acid sequence synthesized by *matR*.

The evolutionary study of the differential loci in the mitogenomes of *T. esculentum* found that the two types of plants fell into two clusters, as expected. The type 1 plants can be further divided into two clades, and the divergence is thought to be possibly related to soil moisture content, although this still needs to be verified by studies with larger sample sizes. Three consecutive substitutions at positions 127,684 to 127,686 on chromosome LS1 (OK638188) may have played an important role here. The phylogenetic tree constructed based on the mitochondrial genes of *T. esculentum* and other related Fabaceae species was consistent with the previously published one built on chloroplast genes. *C. canadensis* and *T. esculentum* were found to be closely related, and they have more complete sets of mitochondrial genes than the other legumes.

Heteroplasmy in the mitochondrial genome of *T. esculentum* is not as prevalent as in its chloroplast genome, but higher levels do exist at certain loci, such as loci 127,684 to 127,686 on chromosome LS1. Among all samples, only one individual A11 from Aminuis had a generally high degree of heteroplasmy at most differential loci, consistent with what was seen in its chloroplast genome. Whether it is because of some of its own characteristics that A11 escaped the genetic bottleneck at the developmental stage is still unclear.

Chloroplast insertions are thought to be concentrated in one of the subgenomic rings of the mitogenome of *T. esculentum*. The mitogenomes of all marama individuals were found to contain a long chloroplast DNA insertion over 9 kb in length. The study of the polymorphisms on this segment infers that the sequence is protected by some mechanism in the chloroplast genome, so only a small amount of synonymous substitutions is retained, but after being inserted into the mitogenome, a large number of mutations have accumulated on it, making the genes on it lose their function.

## Supplementary data

Supplementary materials have been uploaded to JXB including 3 tables and 32 figures.

## Acknowledgments

The authors would like to thank K. Logue for help with the initial genome assembly, to P. Chimwamurombe, M. Takundwa, J. Vorster, and K. Kunert for providing marama samples from Namibia and from the University of Pretoria Farm, to students in BIOL 301/401 in 2015 for assistance with DNA isolation and in BIOL 309 in 2018 for sample collection.

## Author Contributions

J.L. carried out the bioinformatics assembly and data analysis and drafted the manuscript. C.C. conceived initial ideas for the project, provided extracted DNA, and assisted in writing and editing the manuscript. All authors contributed to the submitted article.

## Data availability

The sequencing data used in this study are available and have been deposited under BioProject PRJNA897263 (https://www.ncbi.nlm.nih.gov/bioproject/897263).

