## Supplementary tables and figures for "Comparative Analysis of Tylosema esculentum Mitochondrial DNA Revealed Two Distinct Genome Structures"

Supplementary Materials

Table S1. Summary of WGS Illumina data and sources of the 84 samples

| Sample | Plant source | Raw reads | Raw data | Mito Cvrg | Cytotype |
|---|---|---|---|---|---|
| M17 | NM Farm Seeds | 96750044 | 14512506600 | 2480 | Type 2 |
| S_35 | NM Farm Seeds | 86626894 | 12994034100 | 1928 | Type 2 |
| S_19 | NM Farm Seeds | 55375806 | 8306370900 | 1493 | Type 2 |
| S_4 | NM Farm Seeds | 73435564 | 11015334600 | 2817 | Type 2 |
| S_13 | NM Farm Seeds | 217730028 | 32659504200 | 5368 | Type 2 |
| S_27 | NM Farm Seeds | 107264134 | 16089620100 | 349 | Type 2 |
| S_20 | NM Farm Seeds | 55371622 | 8305743300 | 1455 | Type 2 |
| S_30 | NM Farm Seeds | 106501502 | 15975225300 | 2963 | Type 2 |
| S_33 | NM Farm Seeds | 130811500 | 19621725000 | 4788 | Type 2 |
| M7 | NM Farm Seeds | 172383630 | 25857544500 | 4279 | Type 2 |
| M8 | NM Farm Seeds | 123934796 | 18590219400 | 3713 | Type 2 |
| M1 # | NM Farm Seeds | 170816890 | 25622533500 | 5240 | Type 2 |
| M2 | NM Farm Seeds | 194335620 | 29150343000 | 933 | Type 2 |
| M11 | NM Farm Seeds | 125005082 | 18750762300 | 4213 | Type 2 |
| M12 | NM Farm Seeds | 128360580 | 19254087000 | 1500 | Type 2 |
| M15 | NM Farm Seeds | 134439570 | 20165935500 | 3781 | Type 2 |
| M16 | NM Farm Seeds | 132448260 | 19867239000 | 3789 | Type 2 |
| M23 | NM Farm Seeds | 134475978 | 20171396700 | 3642 | Type 2 |
| M22 | NM Farm Seeds | 111447592 | 16717138800 | 3891 | Type 2 |
| M24 | NM Farm Seeds | 127392478 | 19108871700 | 3546 | Type 2 |
| M26 | NM Farm Seeds | 225925198 | 33888779700 | 699 | Type 2 |
| M28 | NM Farm Seeds | 193804308 | 29070646200 | 4799 | Type 2 |
| M25 | NM Farm Seeds | 145937916 | 21890687400 | 3860 | Type 2 |
| N29 | NM Farm Seeds | 147710832 | 22156624800 | 4863 | Type 2 |
| M31 | NM Farm Seeds | 177899436 | 26684915400 | 3647 | Type 2 |
| M34 | NM Farm Seeds | 124221640 | 18633246000 | 1060 | Type 2 |
| M36 | NM Farm Seeds | 125524008 | 18828601200 | 2242 | Type 2 |
| M37 | NM Farm Seeds | 185151336 | 27772700400 | 6847 | Type 2 |
| M38 | NM Farm Seeds | 168052326 | 25207848900 | 4617 | Type 2 |
| M40 | Namibia Unknown | 135702768 | 20355415200 | 1720 | Type 1 |
| Index1 # | UP Farm | 41499124 | 4149912400 | 269 | Type 2 |
| Index10 | UP Farm Seeds | 36382172 | 3638217200 | 209 | Type 2 |
| Index11 | UP Farm Seeds | 34631932 | 3463193200 | 295 | Type 2 |
| Index12 # | UP Farm | 39339004 | 3933900400 | 804 | Type 2 |
| Index19 # | UP Farm | 35994474 | 3599447400 | 258 | Type 2 |
| Index3 # | UP Farm | 35991990 | 3599199000 | 395 | Type 2 |
| Index5 # | UP Farm | 34349602 | 3434960200 | 814 | Type 2 |

| | | | | | |
|---|---|---|---|---|---|
| Index8 | Namibia Unknown | 42747718 | 4274771800 | 359 | Type 1 |
| Index9 # | UP Farm | 34312880 | 3431288000 | 793 | Type 2 |
| R1R2 | Unknown | 358941018 | 35894101800 | 1550 | Type 2 |
| A1 # | Aminuis Seeds | 93324222 | 13998633300 | 707 | Type 1 |
| A2 | Aminuis Seeds | 93445492 | 14016823800 | 437 | Type 1 |
| A3 | Aminuis Seeds | 84081976 | 12612296400 | 855 | Type 1 |
| A4 | Aminuis Seeds | 60965538 | 9144830700 | 607 | Type 1 |
| A5 | Aminuis Seeds | 91044746 | 13656711900 | 587 | Type 1 |
| A6 | Aminuis Seeds | 89785050 | 13467757500 | 677 | Type 1 |
| A7 | Aminuis Seeds | 86000118 | 12900017700 | 528 | Type 1 |
| A8 | Aminuis Seeds | 57087632 | 8563144800 | 549 | Type 1 |
| A9 # | Aminuis | 62849376 | 9427406400 | 559 | Type 1 |
| A10 # | Aminuis | 75633554 | 11345033100 | 541 | Type 1 |
| A11 # | Aminuis | 92851730 | 13927759500 | 923 | Type 1 |
| A12 # | Aminuis | 84374620 | 12656193000 | 561 | Type 1 |
| A13 # | Aminuis | 83450100 | 12517515000 | 864 | Type 1 |
| nar15 # | Aminuis | 81618952 | 12242842800 | 389 | Type 1 |
| nar16 | UP Farm Seeds | 96595344 | 14489301600 | 807 | Type 2 |
| S1 # | Tsjaka | 10358444 | 1035844400 | 25 | Type 1 |
| S2 # | Tsjaka | 20343934 | 2034393400 | 198 | Type 1 |
| S3 # | Tsjaka | 33100100 | 3310010000 | 597 | Type 1 |
| S4 # | Tsjaka | 25428338 | 2542833800 | 373 | Type 1 |
| S5 # | Okamatapati | 27183834 | 2718383400 | 396 | Type 1 |
| S6 # | Tsumkwe | 24185808 | 2418580800 | 419 | Type 1 |
| S7 # | Tsumkwe | 22075112 | 2207511200 | 162 | Type 1 |
| S8 # | Tsumkwe | 16337544 | 1633754400 | 163 | Type 1 |
| S9 # | Aminuis | 25274640 | 2527464000 | 270 | Type 1 |
| S10 # | Aminuis | 28329944 | 2832994400 | 311 | Type 1 |
| S11 # | Aminuis | 26741342 | 2674134200 | 288 | Type 1 |
| S12 # | Aminuis | 29520092 | 2952009200 | 423 | Type 1 |
| S13 # | Aminuis | NA | NA | 236 | Type 1 |
| S14 # | Aminuis | 22928100 | 2292810000 | 283 | Type 1 |
| S15 # | Aminuis | 22856344 | 2285634400 | 486 | Type 1 |
| S16 # | Aminuis | 25333752 | 2533375200 | 338 | Type 1 |
| S17 # | Aminuis | 23418786 | 2341878600 | 258 | Type 1 |
| S18 # | Tsumkwe | 25764102 | 2576410200 | 234 | Type 1 |
| S19 # | Aminuis | 24407080 | 2440708000 | 421 | Type 1 |
| S20 # | Osire | 25692398 | 2569239800 | 407 | Type 1 |
| S21 # | Osire | 25940358 | 2594035800 | 381 | Type 1 |
| S22 # | Osire | 32107310 | 3210731000 | 609 | Type 1 |
| S23 # | Osire | 35844166 | 3584416600 | 697 | Type 1 |
| S24 # | Tsumkwe | 31973008 | 3197300800 | 411 | Type 1 |
| S25 # | Ombujondjou | 24401422 | 2440142200 | 539 | Type 1 |

| | | | | | |
|---|---|---|---|---|---|
| S26 # | Ombujondjou | 32758062 | 3275806200 | 433 | Type 1 |
| S27 # | Epukiro | 28122544 | 2812254400 | 573 | Type 1 |
| S28 # | Epukiro | 31273940 | 3127394000 | 699 | Type 1 |
| S29 # | Otjiwarongo | 25274640 | 2527464000 | 498 | Type 1 |

**#** 43 independent samples for sequence diversity and phylogenetic analysis; Mito Cvrg = approximate read depth for single-copy regions of the mitogenome

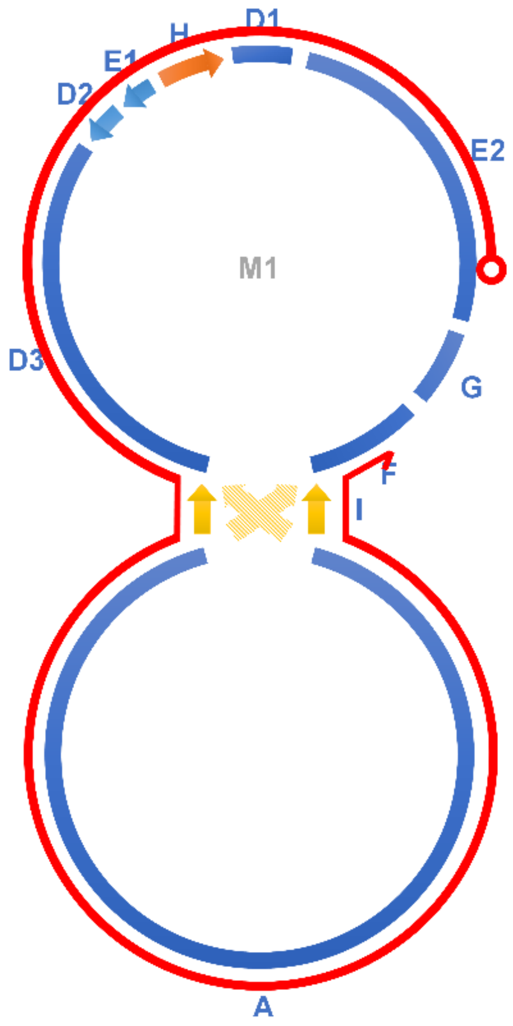

Figure S1. Alignment of contig tig00000040 of 159,605 bp, assembled by HiCanu on the Sample 4 PacBio HiFi reads, to the type 2 marama subgenomic chromosome M1. The Canu input genome size was set to 2M to better capture complete organelle genome sequences. The red curve indicates where the contig is aligned, starting from the red circle and ending with the red arrow.

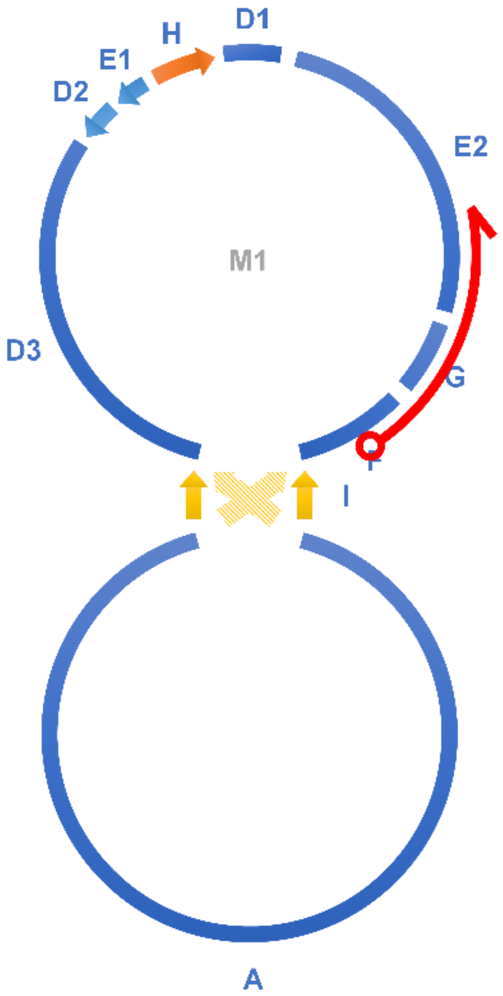


Figure S2. Alignment of contig tig00000050 of 13,828 bp, assembled by HiCanu on the Sample 4 PacBio HiFi reads, to the type 2 marama subgenomic chromosome M1. The Canu input genome size was set to 2M to better capture complete organelle genome sequences. The red curve indicates where the contig is aligned, starting from the red circle and ending with the red arrow. This together with tig00000040 (Figure S1) verifies the structure of M1. Discontinuous assembly results from fluctuating in coverage at some locations, and the complete structure has been verified by manual extension.

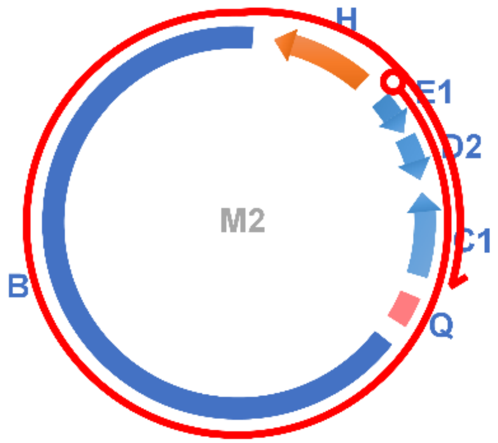


Figure S3. Alignment of contig tig00000055 of 64,325 bp, assembled by HiCanu on the Sample 4 PacBio HiFi reads, to the type 2 marama subgenomic chromosome M2. The Canu input genome size was set to 2M to better capture complete organelle genome sequences. The red curve indicates where the contig is aligned, starting from the red circle and ending with the red arrow. This contig validates the circular structure of M2.

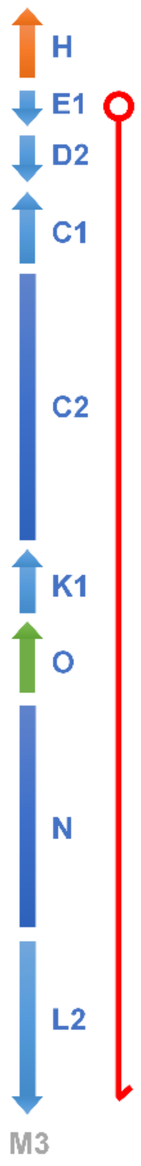


Figure S4. Alignment of contig tig00000057 of 87,648 bp, assembled by HiCanu on the Sample 4 PacBio HiFi reads, to the type 2 marama subgenomic chromosome M3. The Canu input genome size was set to 2M to better capture complete organelle genome sequences. The red line indicates where the contig is aligned, starting from the red circle and ending with the red arrow.

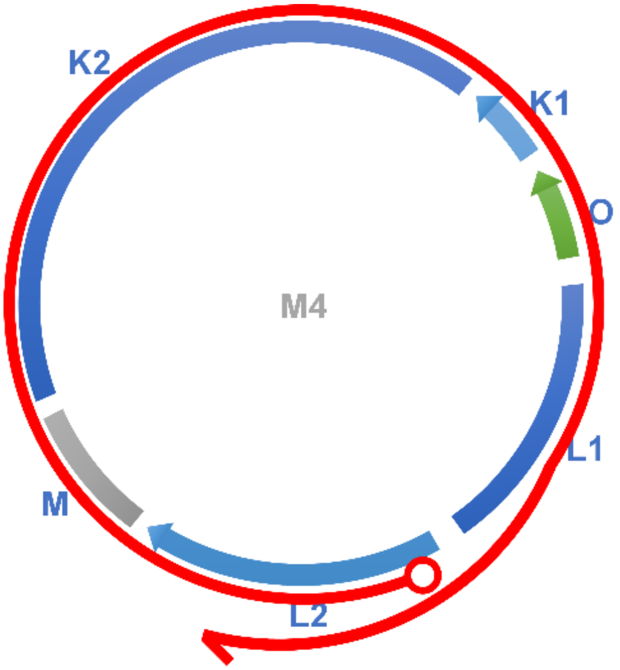


Figure S5. Alignment of contig tig00000058 of 123,793 bp, assembled by HiCanu on the Sample 4 PacBio HiFi reads, to the type 2 marama subgenomic chromosome M4. The Canu input genome size was set to 2M to better capture complete organelle genome sequences. The red curve indicates where the contig is aligned, starting from the red circle and ending with the red arrow. This verifies the circular structure of M4.

Two different connections were seen at the pair of inverted repeats I, which formed a normal circular molecule, or an 8-shaped ring resulting from recombination on the repeats ($A^{1}$-I-D3 and $A^{82,874}$-I-F became $A^{82,874}$-I-D3 and $A^{1}$-I-F). Both molecules were confirmed by the PacBio HiFi reads of Sample 4 and found to be in very close proportions, as shown in Figure S6-9.

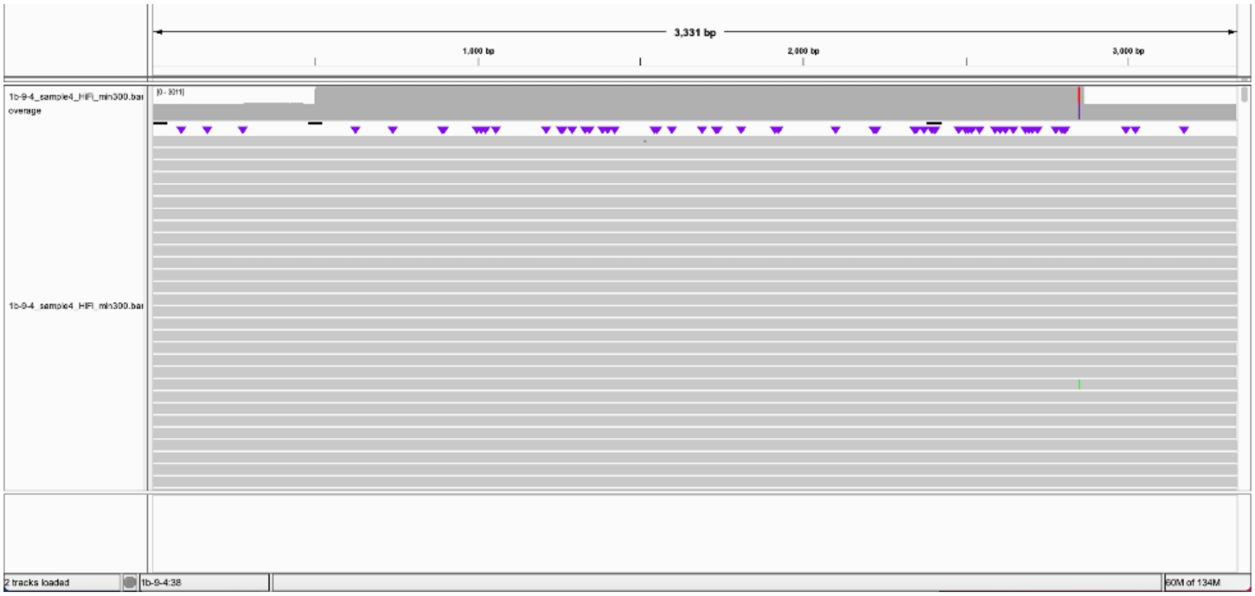


Figure S6. Alignment of Sample 4 PacBio HiFi reads to the artificial chromosome concatenated by node A (82,375 - 82,874 bp), node I, and node D3 (26,671-26,172 bp) using pbmm2. The minimum length was set to 300 to avoid interference from homologous DNA fragments. The coverage of the inverted repeat node I was doubled as expected.

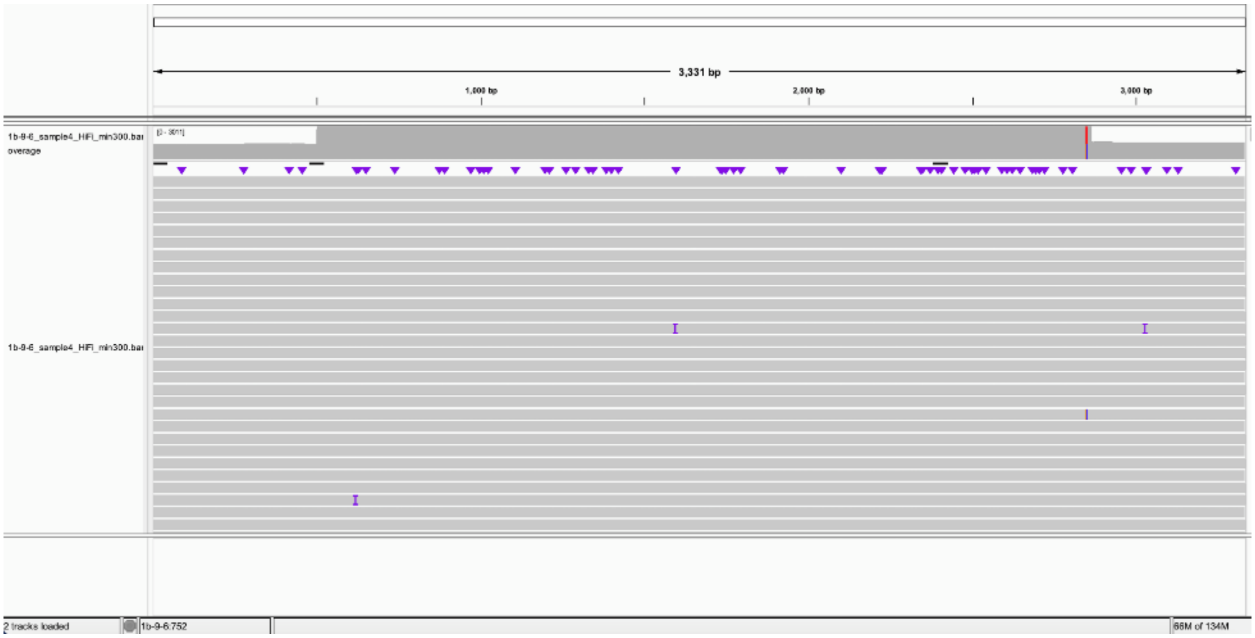


Figure S7. Alignment of Sample 4 PacBio HiFi reads to the artificial chromosome concatenated by node A (82,375 - 82,874 bp), node I, and node F (8,590 – 8,091 bp) using pbmm2. The minimum length was set to 300 to avoid interference from homologous DNA fragments. The coverage of the inverted repeat node I was doubled as expected.

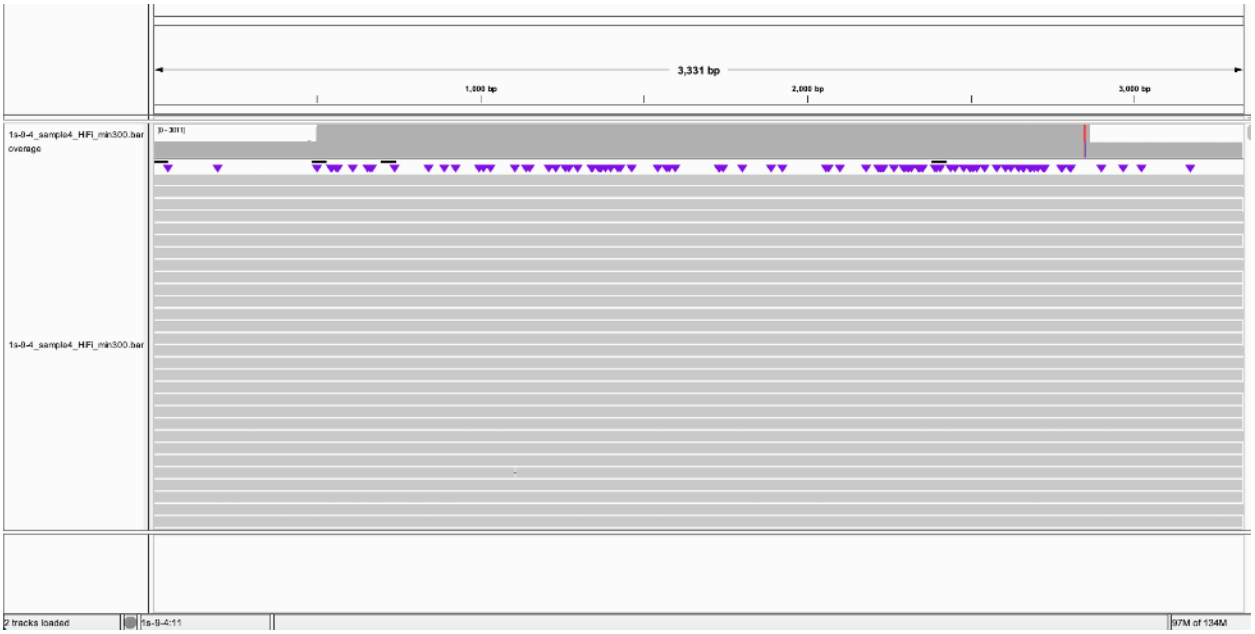


Figure S8. Alignment of Sample 4 PacBio HiFi reads to the artificial chromosome concatenated by node A (500 - 1 bp), node I, and node D3 (26,671-26,172 bp) using pbmm2. The minimum length was set to 300 to avoid interference from homologous DNA fragments. The coverage of the inverted repeat node I was doubled as expected.

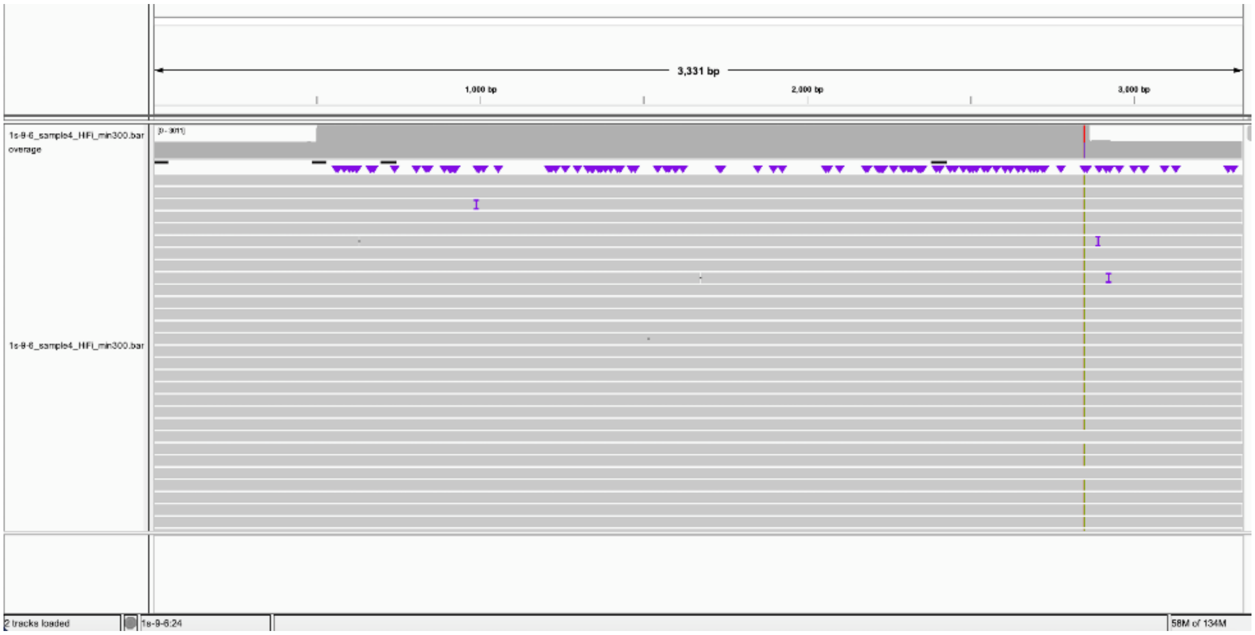


Figure S9. Alignment of Sample 4 PacBio HiFi reads to the artificial chromosome concatenated by node A (500 - 1 bp), node I, and node F (8,590 – 8,091 bp) using pbmm2. The minimum length was set to 300 to avoid interference from homologous DNA fragments. The coverage of the inverted repeat node I was doubled as expected.

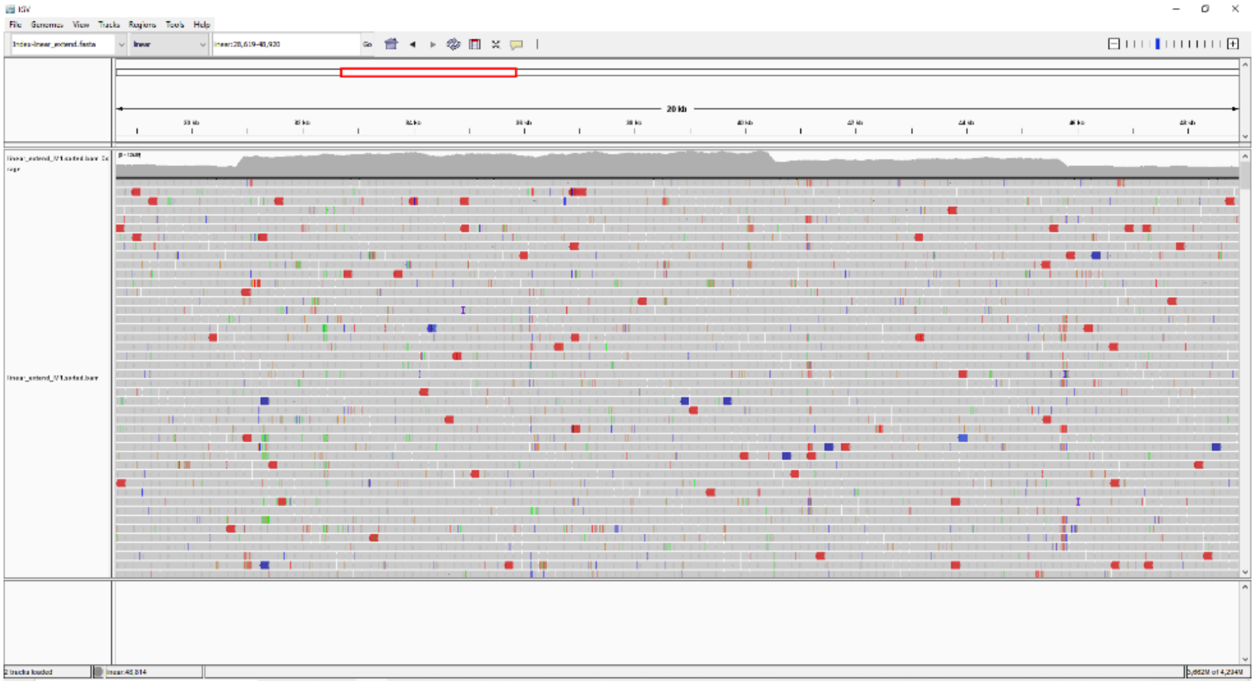

Figure S10. IGV visualization of WGS Illumina reads of individual M1 aligned to the one end of the linear chromosome M3 (extended by the sequence of D1, thus arranged as D1-H-E1-D2-C1-C2). A clear increase of coverage from one copy to two, and then to three can be seen from right to left. The two ends, C2 and D1 have only one copy. C1 has two copies, as it is owned by both chromosomes M2 and M3. H-E1-D2 is a long repeat present in three chromosomes, M1, M2, and M3, with tripled depth as shown.

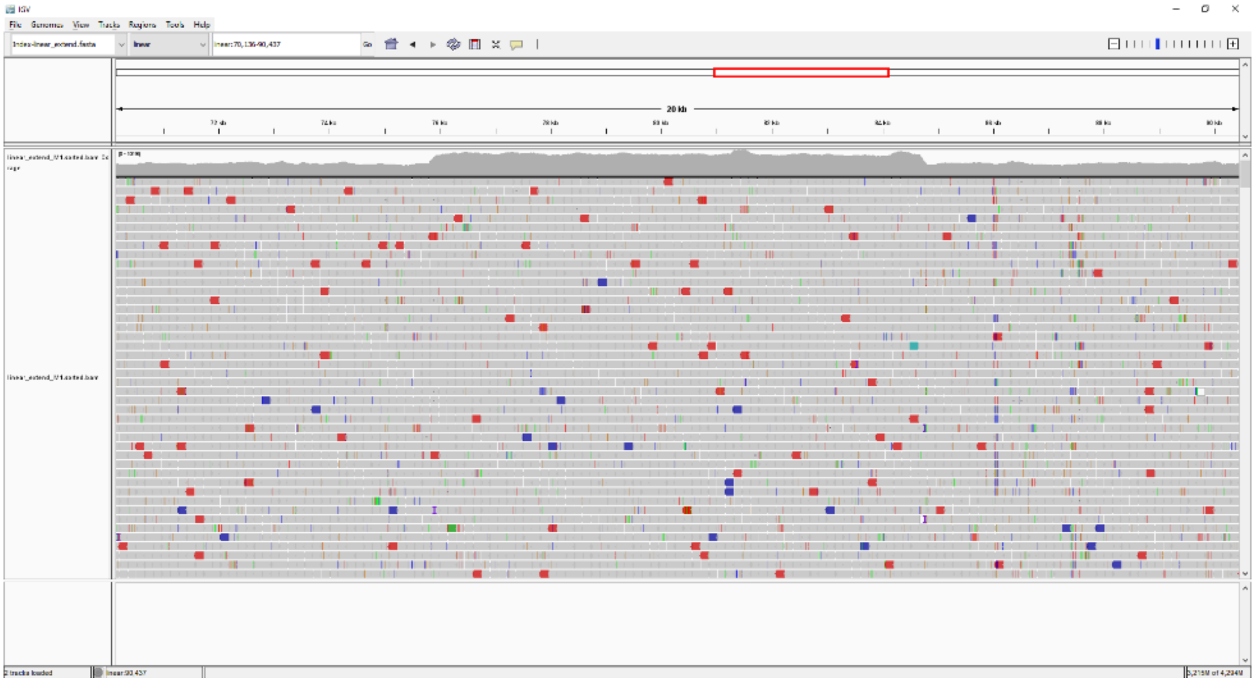

Figure S11. IGV visualization of WGS Illumina reads of individual M1 aligned to the mitochondrial genome fragment C2-K1-O-N via Bowtie2. The coverage of K1 and O in the middle is doubled because it is a long repeat owned by two chromosomes, M3 and M4.

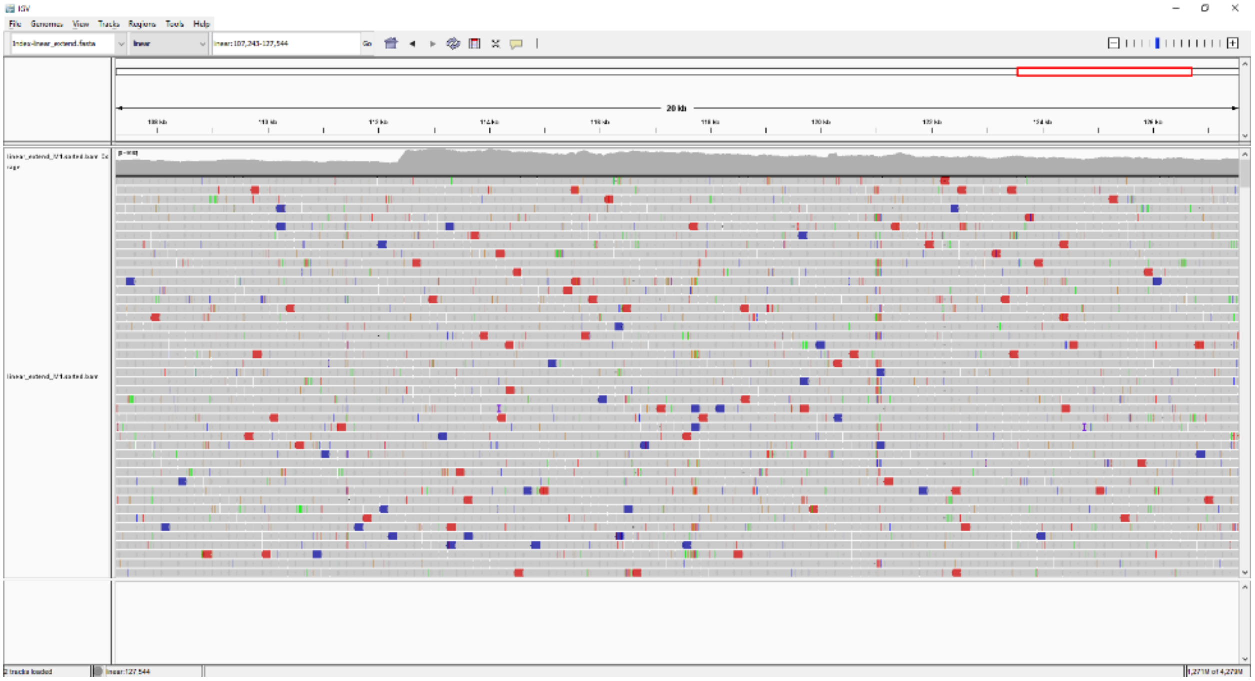

Figure S12. IGV visualization of WGS Illumina reads of marama individual M1 aligned to the mitochondrial genome fragment L1-L2 by Bowtie2. From L2 onwards, the read depth is doubled because L2 is a segment owned by both chromosomes M3 and M4. However, the coverage gradually decreases from left to right after doubling, as the end of the linear chromosome M3 is reached.

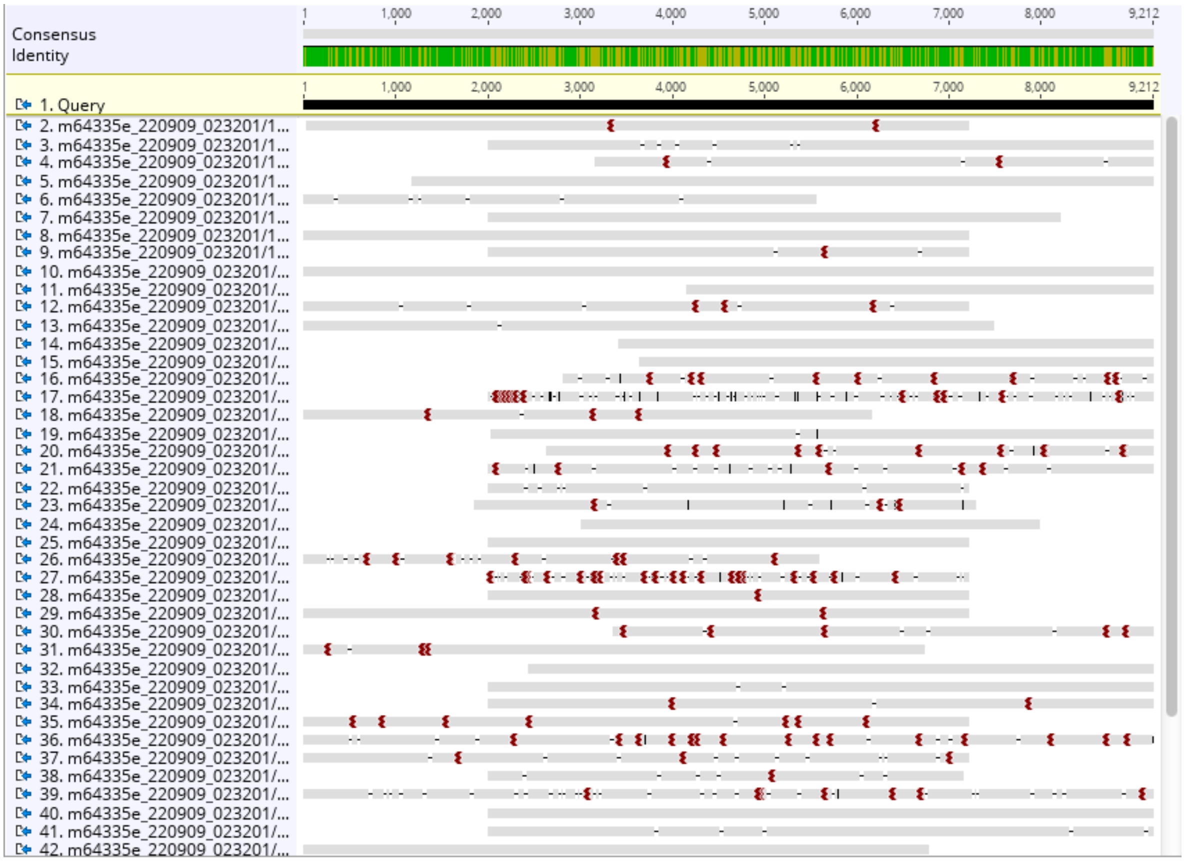


Figure S13. The PacBio HiFi reads of the type 1 individual Sample 32 were mapped to the fragment B-H-B (2000 bp sequences were taken from both ends of node B, with node H in the middle) by BLAST in Geneious 9. Reads going across the entire fragment B-H-B can be seen, indicating the presence of a closed ring consisting only of nodes B and H in type 1 individual Sample 32. In addition, reads from chromosome M1, mapped only to node H, were also observed. Reads, mapped to node H and one end of node B (the other end entered chromosome M1, not shown here), were seen, representing a combination of the two subgenomic structures.

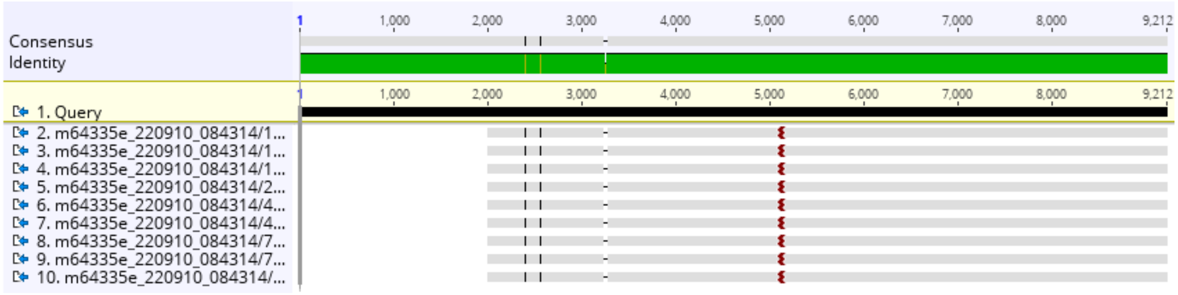


Figure S14. The PacBio HiFi reads of the type 2 individual Sample 4 were mapped to the fragment B-H-B (2000 bp sequences were taken from both ends of node B, with node H in the middle) by BLAST in Geneious 9. Node H was only connected to one end of node B but not to the other. The closed ring consisting only of node B and H does not exist in type 2 individual Sample 4.

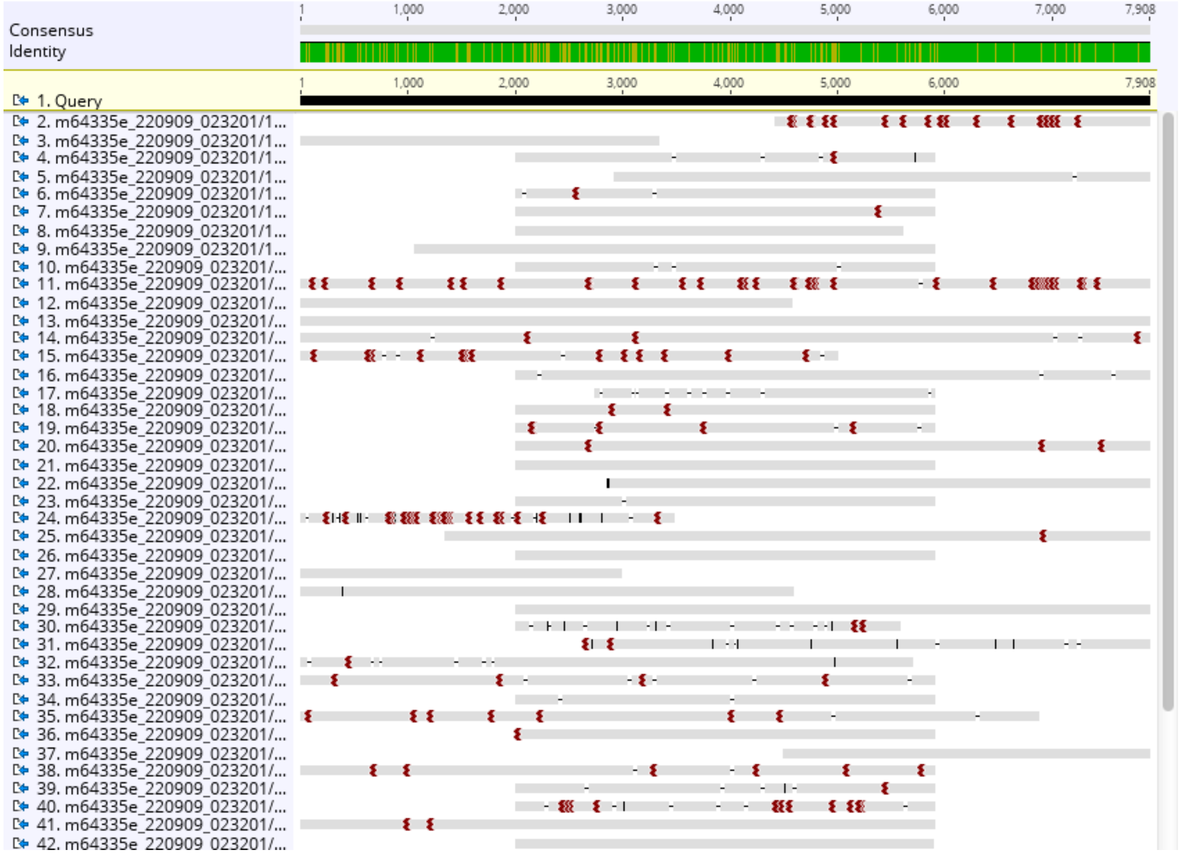


Figure S15. The PacBio HiFi reads of the type 1 individual Sample 32 were mapped to the fragment C-A-C (node A, 39,444-43,351 bp, surrounded by 2000 bp sequences at both ends of node C) by BLAST in Geneious 9. Reads spanning the entire fragment C-A-C can be seen, indicating the existence of a closed subgenomic ring consisting only of nodes C and A in type 1 individual Sample 32. Furthermore, reads from chromosome M1 were observed to map only to node A in the middle, and also reads were seen to map to node A and one end of node C (the other end entered chromosome M1, not shown here), representing a combination of the two subgenomic structures.

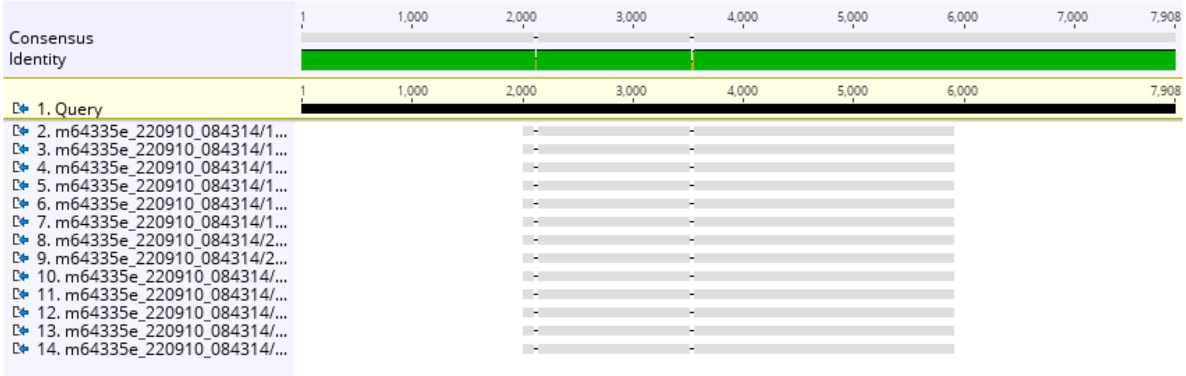

Figure S16. The PacBio HiFi reads of the type 2 individual Sample 4 were mapped to the fragment C-A-C (node A, 39,444-43,351 bp, surrounded by 2000 bp sequences at both ends of node C) by BLAST in Geneious 9. Node A (39,444-43,351 bp) was not connected to either end of node C in Sample 4, suggesting that the two types of mitochondrial genomes have distinct structures in this region.

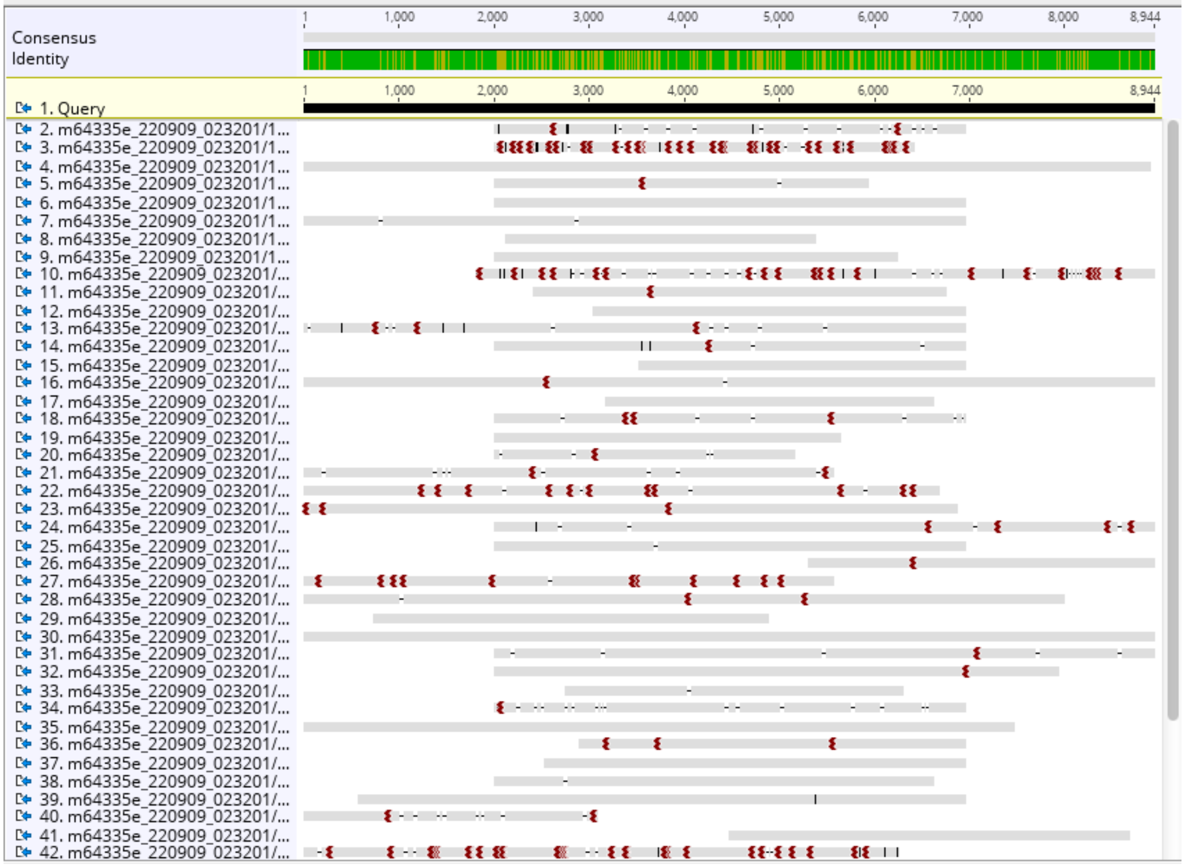


Figure S17. The PacBio HiFi reads of the type 1 individual Sample 32 were mapped to the fragment N-O-N (2000 bp sequences were taken from both ends of node N, with node O in the middle) by BLAST in Geneious 9. Reads spanning the entire fragment N-O-N can be seen, indicating the existence of a closed subgenomic ring consisting only of nodes N and O in type 1 individual Sample 32. Furthermore, reads from chromosome M1 were observed to map only to node O in the middle, and also reads were seen to map to node O and one end of node N (the other end entered chromosome M1, not shown here), representing a combination of the two subgenomic structures.

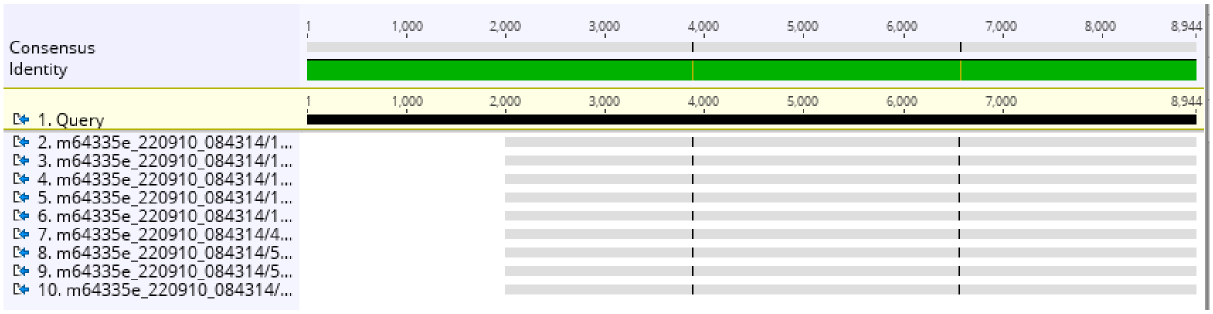


Figure S18. The PacBio HiFi reads of the type 2 individual Sample 4 were mapped to the fragment N-O-N (2000 bp sequences were taken from both ends of node N, with node O in the middle) by BLAST in Geneious 9. Node O was only connected to one end of node N but not to the other. The closed ring consisting only of node N and O does not exist in type 2 individual Sample 4.

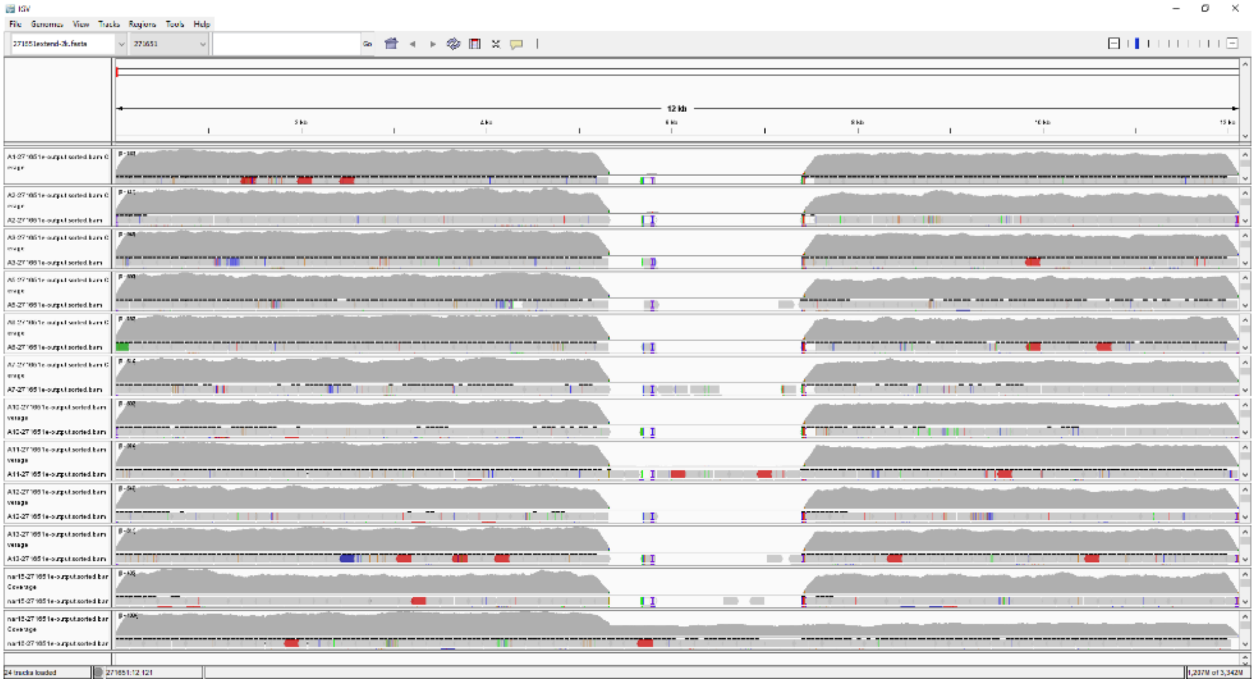

Figure S19. Bowtie 2 alignment of WGS Illumina reads from 12 marama samples to the 12,113 bp fragment from the type 2 mitochondrial genome chromosome M2, visualized in IGV. The reference sequence starts at C1 and ends at B (Figure 1), with the 2,108 bp type 2 exclusive sequence in between, from 5,325 bp to 7,432 bp. The 10 A plants and nar15, all originally from Aminuis, do not contain this 2,108 bp fragment, indicating that they have a type 1 mitochondrial genome. Furthermore, in these plants, the coverage of C1 is close to that of B, as expected for the type 1 structure. nar16 is a descendent of plants grown on Pretoria Farm and has a type 2 mitochondrial genome. Not only does it contain this 2,108 bp fragment, but it also has a depth-doubled C1 because that’s a repeat sequence that both M2 and M3 have, as shown in Figure 1.

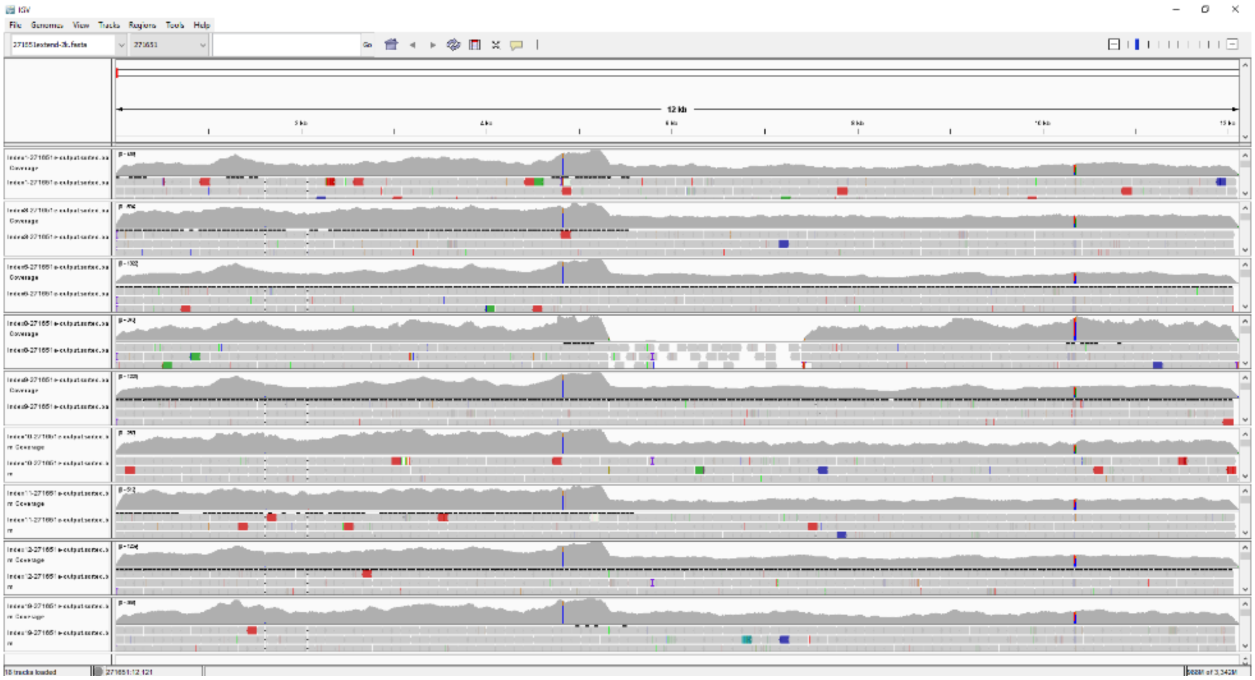

Figure S20. Bowtie 2 alignment of WGS Illumina reads from 9 Index plants to the 12,113 bp fragment from the type 2 mitochondrial genome chromosome M2, visualized in IGV. The reference sequence starts at C1 and ends at B (Figure 1), with the 2,108 bp type 2 specific sequence in between, from 5,325 bp to 7,432 bp. Among these plants, Index8, as the only one not originating from the Pretoria Farm, does not contain this 2,108 bp fragment, indicating that it has a type 1 mitochondrial genome. All remaining Index plants were either collected from the Pretoria Farm or were grown from seeds collected there. They all have type 2 mitochondrial genomes and they all contain this 2,108 bp fragment. Besides, the depth of C1 is doubled in all these plants except Index8 because C1 is a repeat sequence that both M2 and M3 have in the type 2 mitogenome, as shown in Figure 1.

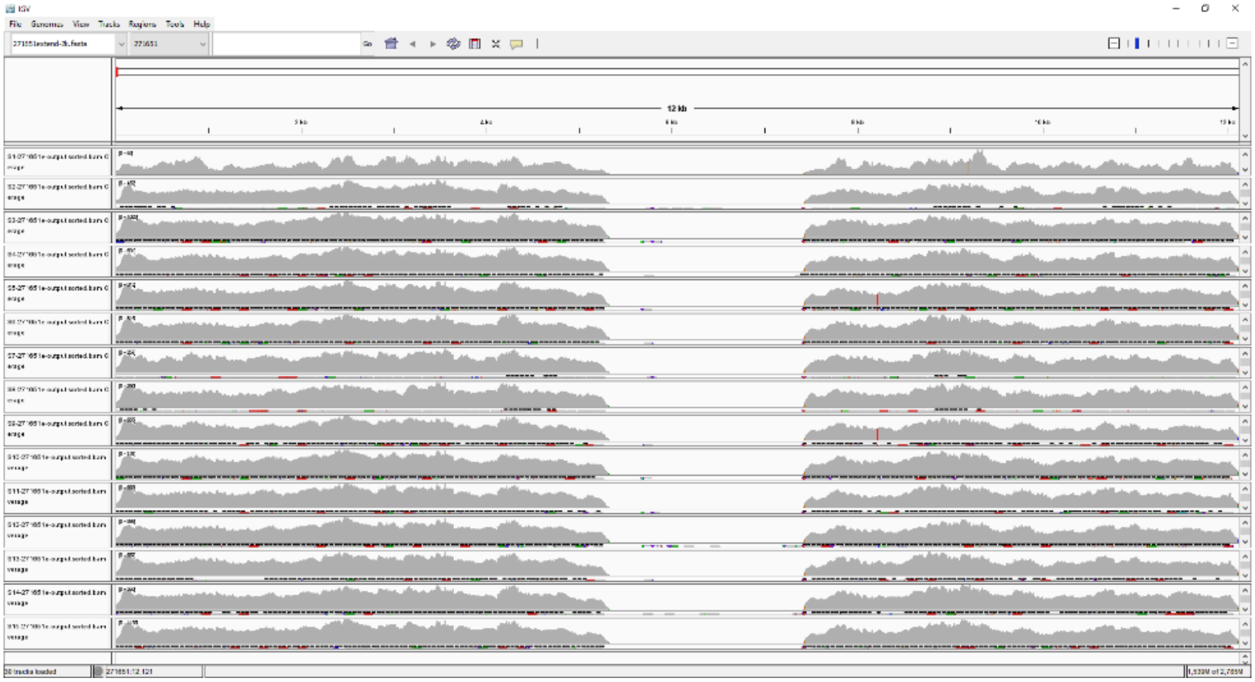

Figure S21. Bowtie 2 alignment of WGS Illumina reads from 15 S plants (S1-S15) to the 12,113 bp fragment from the type 2 mitochondrial genome chromosome M2, visualized in IGV. The reference sequence starts at C1 and ends at B (Figure 1), with the 2,108 bp type 2 specific sequence in between, from 5,325 bp to 7,432 bp. S plants are wild plants collected from 8 different geographical locations in Namibia. None of them contain this 2,108 bp fragment, indicating that they all have the type 1 mitochondrial genome.

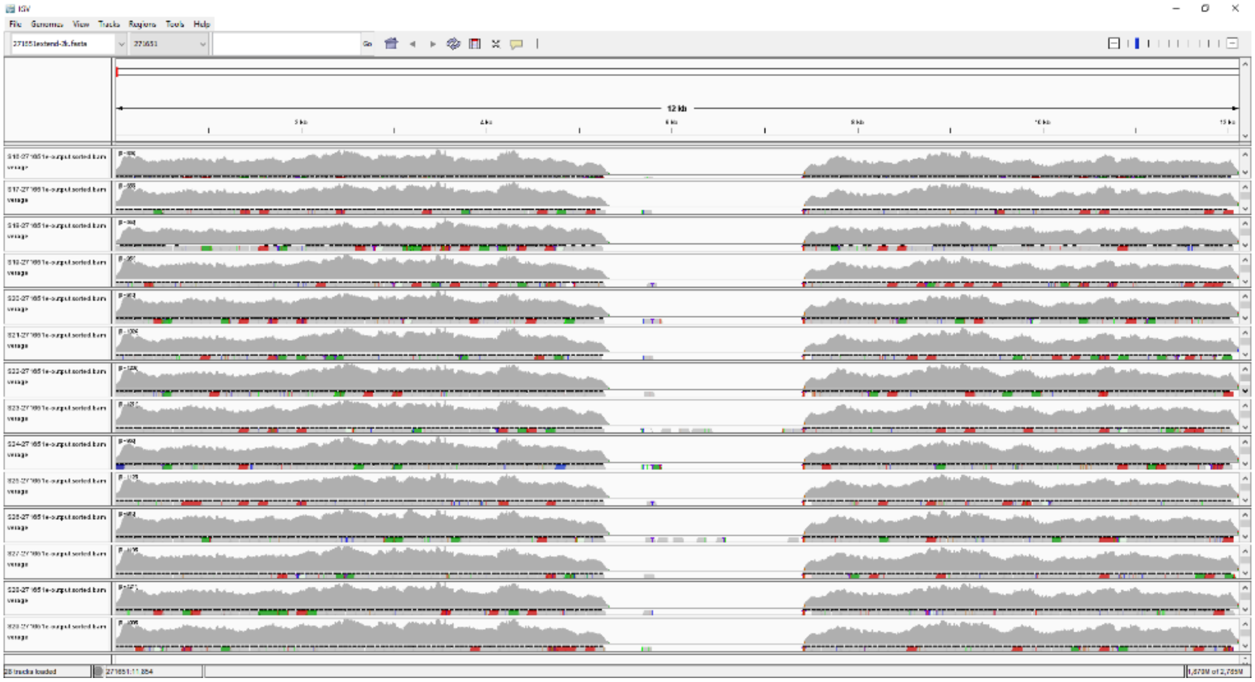

Figure S22. Bowtie 2 alignment of WGS Illumina reads from 14 S plants (S16-S29) to the 12,113 bp fragment from the type 2 mitochondrial genome chromosome M2, visualized in IGV. The reference sequence starts at C1 and ends at B (Figure 1), with the 2,108 bp type 2 specific sequence in between, from 5,325 bp to 7,432 bp. S plants are wild plants collected from 8 different geographical locations in Namibia. None of them contain this 2,108 bp fragment, indicating that they all have the type 1 mitochondrial genome.

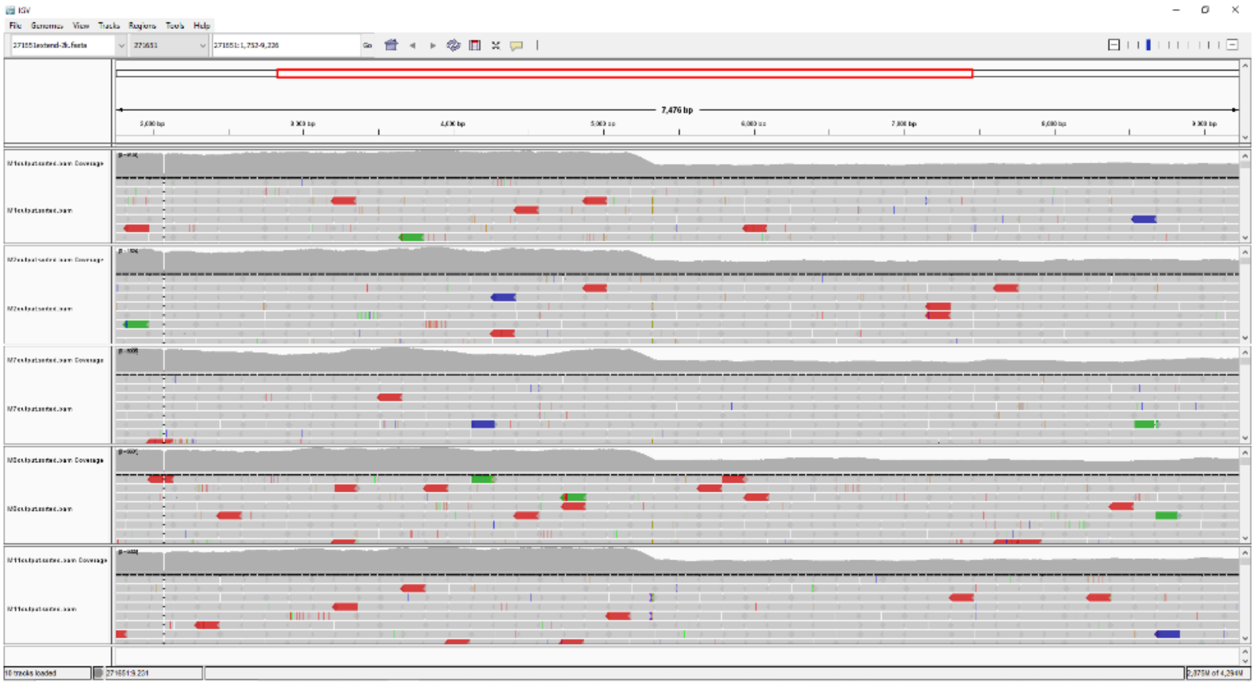

Figure S23. Bowtie 2 alignment of WGS Illumina reads from 5 M plants (M1, M2, M7, M8, and M11) to the 12,113 bp fragment from the type 2 mitochondrial genome chromosome M2, visualized in IGV. The reference sequence starts at C1 and ends at B (Figure 1), with the 2,108 bp type 2 specific sequence in between, from 5,325 bp to 7,432 bp. M plants (except M40) were grown from seeds collected from Namibia Farm. They all contain this 2,108 bp fragment, indicating that they all have the type 2 mitochondrial genome. Furthermore, the coverage of the C1 region is doubled in these plants because it is a repeat sequence that both chromosomes M2 and M3 have in the type 2 mitogenome, as shown in Figure 1.

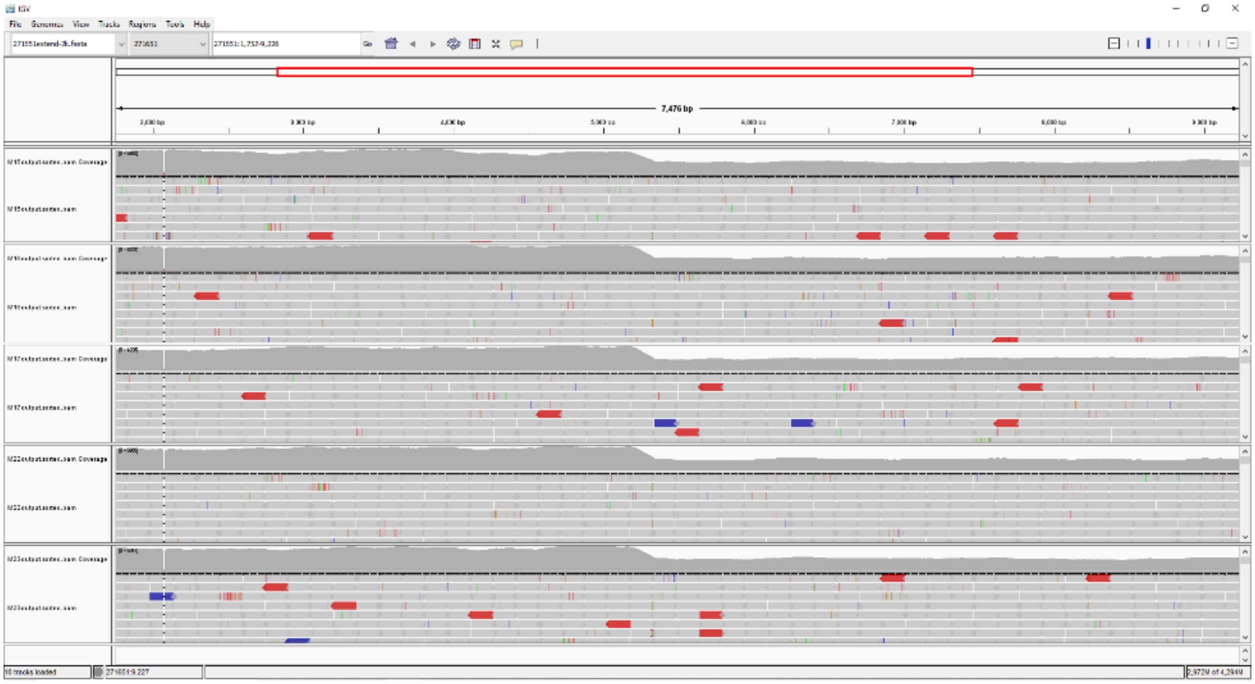

Figure S24. Bowtie 2 alignment of WGS Illumina reads from 5 M plants (M15, M16, M17, M22, and M23) to the 12,113 bp fragment from the type 2 mitochondrial genome chromosome M2, visualized in IGV. The reference sequence starts at C1 and ends at B (Figure 1), with the 2,108 bp type 2 specific sequence in between, from 5,325 bp to 7,432 bp. M plants (except M40) were grown from seeds collected from Namibia Farm. They all contain this 2,108 bp fragment, indicating that they all have type 2 mitochondrial genomes. Furthermore, the coverage of the C1 region is doubled in these 5 plants because it is a repeat sequence that both chromosomes M2 and M3 have in the type 2 mitogenome, as shown in Figure 1.

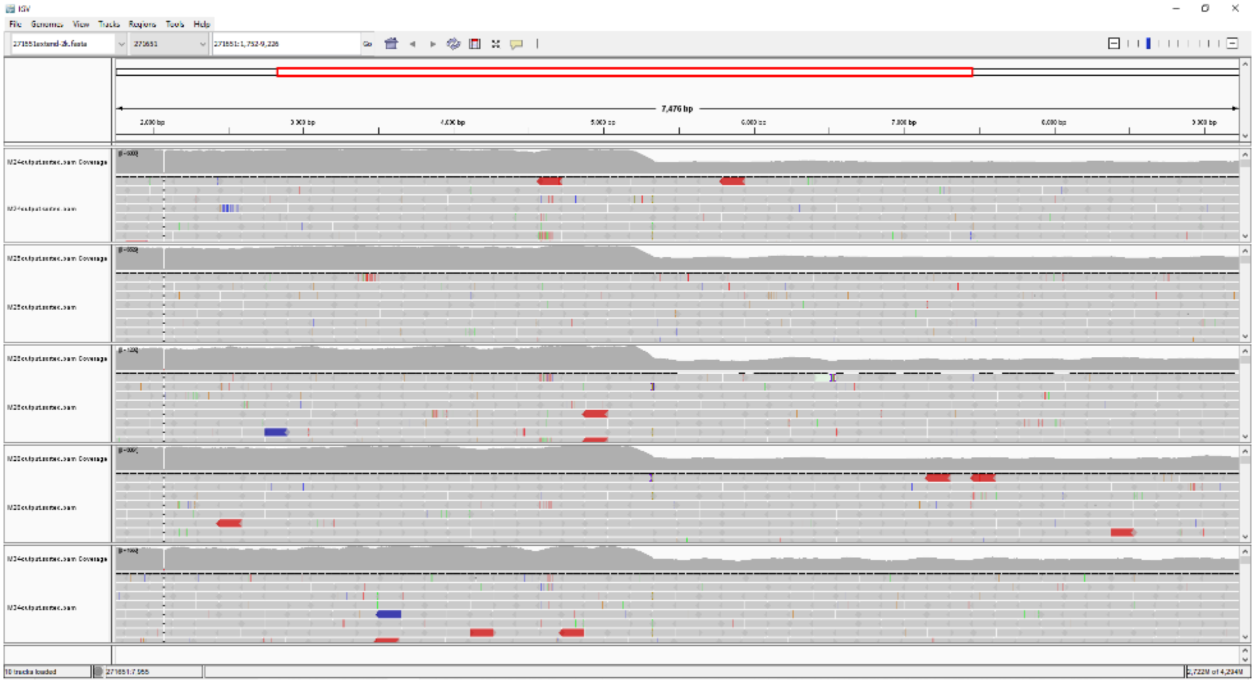

Figure S25. Bowtie 2 alignment of WGS Illumina reads from 5 M plants (M24, M25, M26, M28, and M34) to the 12,113 bp fragment from the type 2 mitochondrial genome chromosome M2, visualized in IGV. The reference sequence starts at C1 and ends at B (Figure 1), with the 2,108 bp type 2 specific sequence in between, from 5,325 bp to 7,432 bp. M plants (except M40) were grown from seeds collected from Namibia Farm. They all contain this 2,108 bp fragment, indicating that they all have type 2 mitochondrial genomes. Furthermore, the coverage of the C1 region is doubled in these plants because it is a repeat sequence that both chromosomes M2 and M3 have in the type 2 mitogenome, as shown in Figure 1.

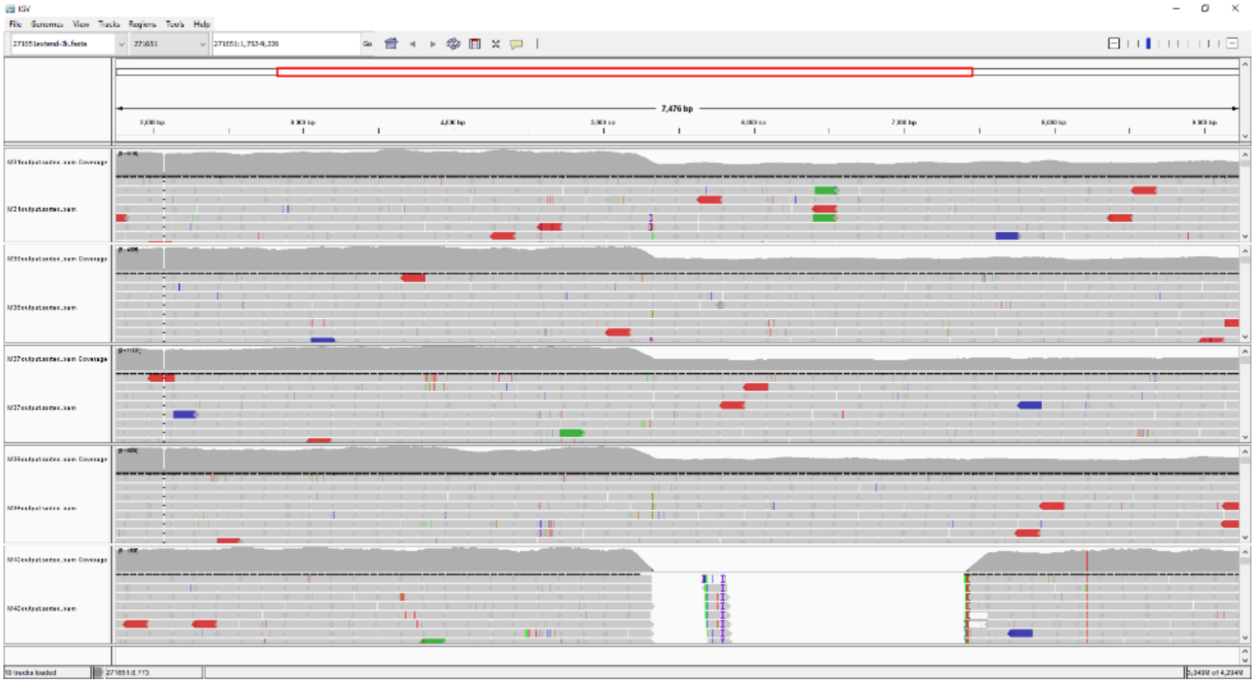

Figure S26. Bowtie 2 alignment of WGS Illumina reads from 5 M plants (M31, M36, M37, M38, and M40) to the 12,113 bp fragment from the type 2 mitochondrial genome chromosome M2, visualized in IGV. The reference sequence starts at C1 and ends at B (Figure 1), with the 2,108 bp type 2 specific sequence in between, from 5,325 bp to 7,432 bp. All M plants (except M40) were grown from seeds collected from Namibia Farm. The origin of sample M40 is unknown. All plants except M40 contain this 2,108 bp fragment, indicating that they have type 2 mitochondrial genomes. Furthermore, the coverage of the C1 region is doubled in these 4 plants because it is a repeat sequence that both chromosomes M2 and M3 have in the type 2 mitogenome, as shown in Figure 1. M40 doesn't have this 2,108 bp fragment, and its C1 and B have close sequencing coverage, suggesting that M40 has a type 1 mitogenome.

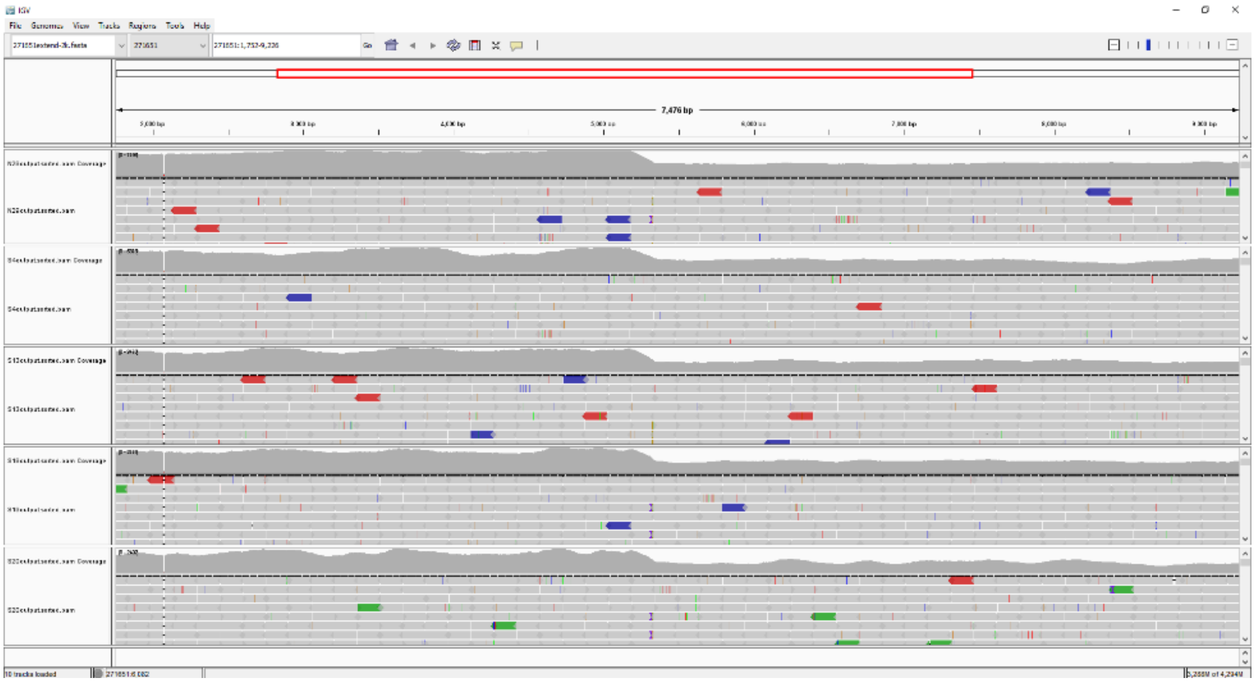

Figure S27. Bowtie 2 alignment of WGS Illumina reads from 5 M individuals (N29, S_4, S_13, S_19, and S_20) to the 12,113 bp fragment from the type 2 mitochondrial genome chromosome M2, visualized in IGV. The reference sequence starts at C1 and ends at B (Figure 1), with the 2,108 bp type 2 specific sequence in between, from 5,325 bp to 7,432 bp. M plants (except M40) were grown from seeds collected from Namibia Farm. They all contain this 2,108 bp fragment, indicating that they all have type 2 mitochondrial genomes. Furthermore, the coverage of the C1 region is doubled in these plants because it is a repeat sequence that both chromosomes M2 and M3 have in the type 2 mitogenome, as shown in Figure 1.

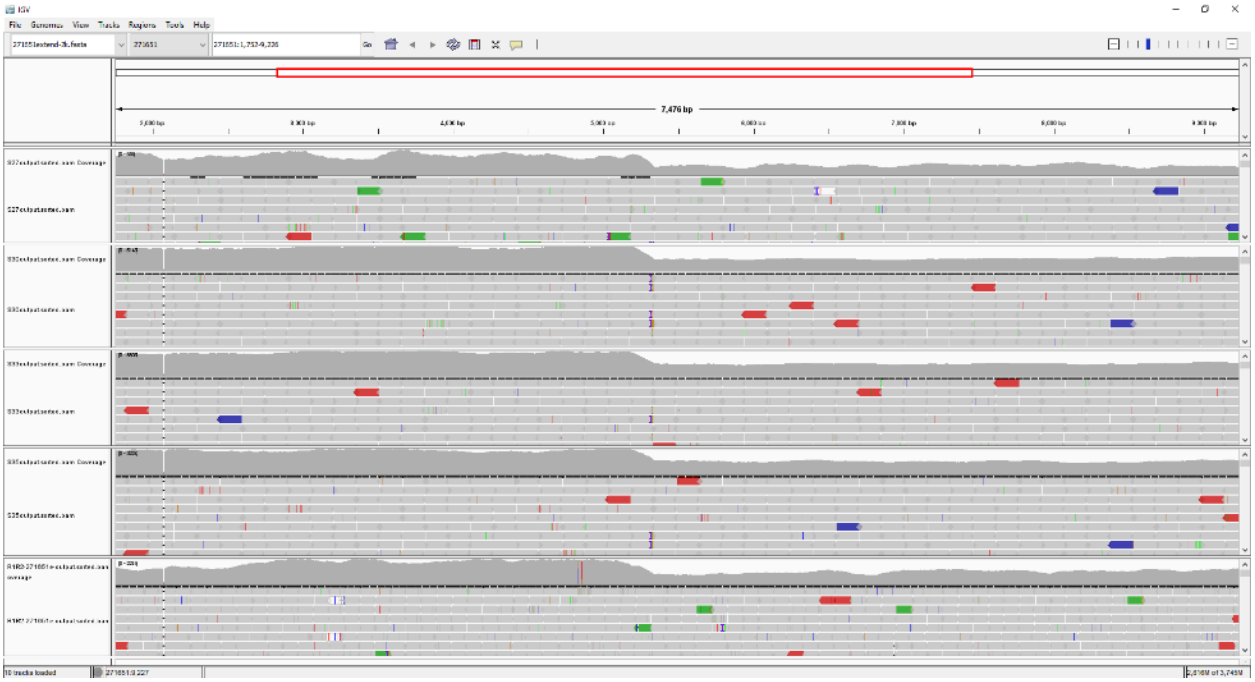

Figure S28. Bowtie 2 alignment of WGS Illumina reads from 4 M plants (S_27, S_30, S_33, and S_35) and R1R2 to the 12,113 bp fragment from the type 2 mitochondrial genome chromosome M2, visualized in IGV. The reference sequence starts at C1 and ends at B (Figure 1), with the 2,108 bp type 2 specific sequence in between, from 5,325 bp to 7,432 bp. M plants (except M40) were grown from seeds collected from Namibia Farm. The origin of sample R1R2 is unknown. However, they all contain this 2,108 bp fragment, indicating that they all have type 2 mitochondrial genomes. Furthermore, the coverage of the C1 region is doubled in all these 5 individuals, which consistent with the type 2 mitogenome structure, as C1 is a repeat sequence contained in both chromosomes M2 and M3 in the type 2 mitogenome, as shown in Figure 1.

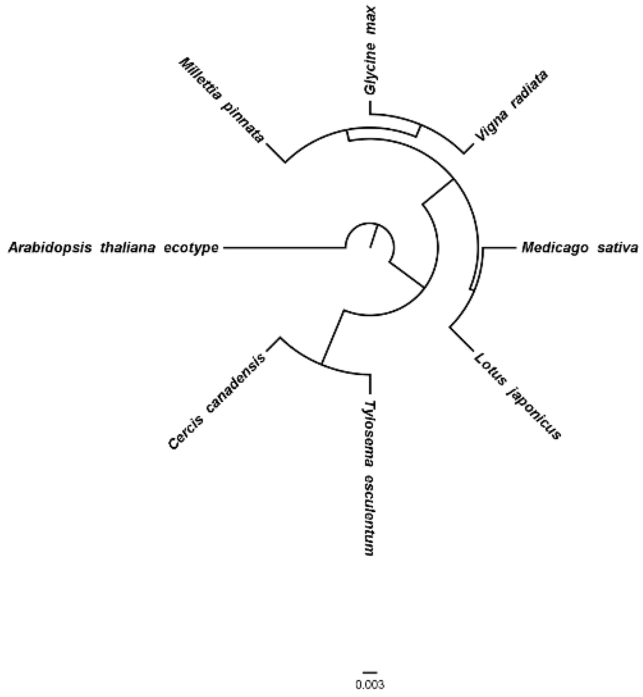


Figure 29. Bayesian inference tree on artificial chromosomes concatenated by 24 conserved mitochondrial genes, *atp1*, *atp4*, *atp6*, *atp8*, *atp9*, *nad3*, *nad4*, *nad4L*, *nad6*, *nad7*, *nad9*, *mttB*, *matR*, *cox1*, *cox3*, *cob*, *ccmFn*, *ccmFc*, *ccmC*, *ccmB*, *rps3*, *rps4*, *rps12*, and *rpl16* from the mitochondrial genomes of *Arabidopsis thaliana* (NC_037304.1), *Cercis canadensis* (MN017226.1), *Lotus japonicus* (NC_016743.2), *Medicago sativa* (ON782580.1), *Millettia pinnata* (NC_016742.1), *Glycine max* (NC_020455.1), and *Vigna radiata* (NC_015121.1) in NCBI drawn by BEAST and FigTree.

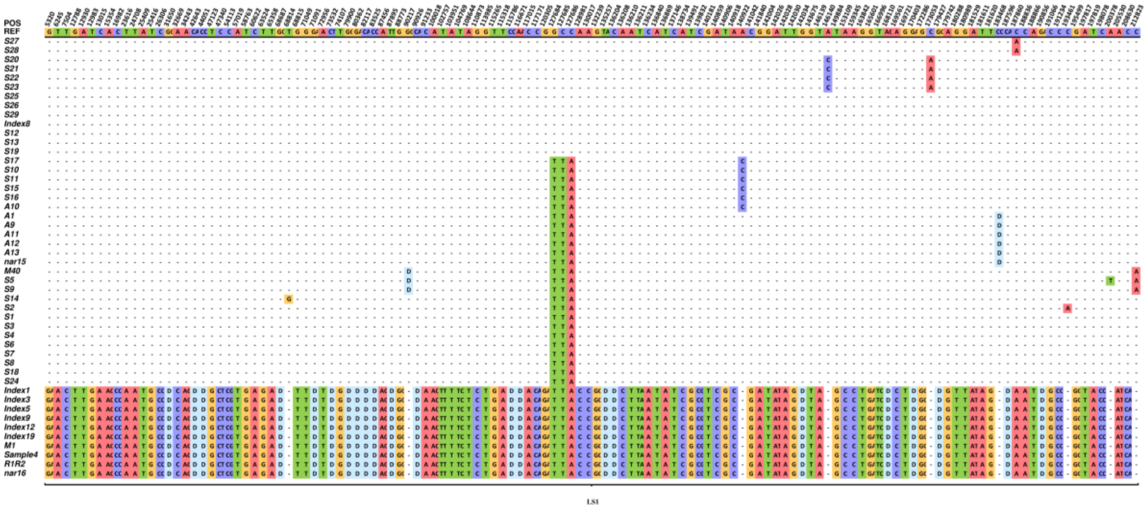


Figure S30. Distribution of variations discovered when aligning the WGS reads of 38 type 1 individuals and 10 type 2 plants to the first 212,823 bp of the marama reference mitogenome chromosome LS1, OK638188. The first row shows the bases in the reference sequence, from the second row onwards only the bases that differ from the reference are shown, and the bases that are the same as the reference are replaced by dashes. All insertions are represented by the first two bases and deletions by the letter “D”.

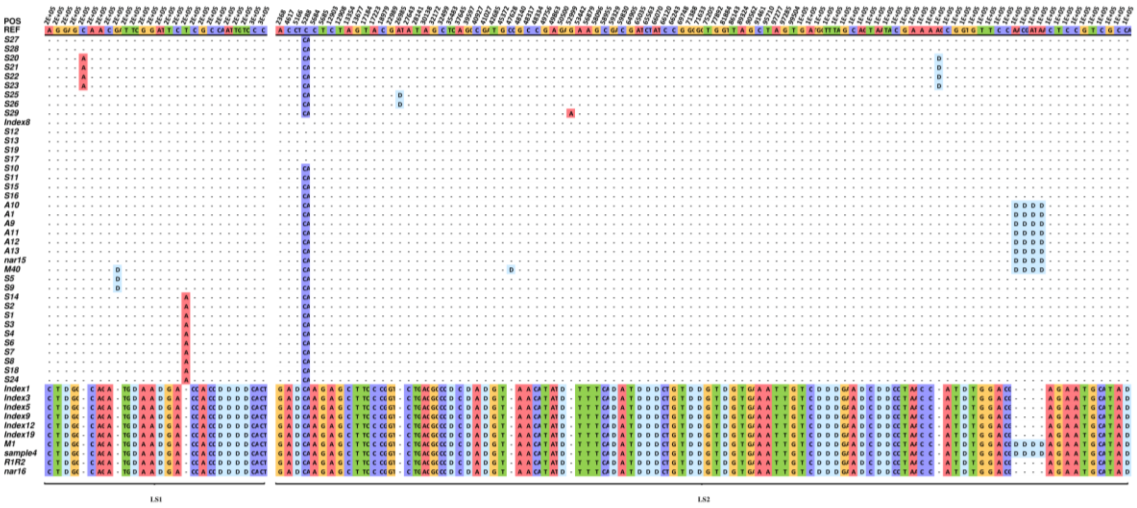


Figure S31. Distribution of variations discovered when aligning the WGS reads of 38 type 1 individuals and 10 type 2 plants to the marama reference mitogenome chromosomes LS1 (OK638188, 212,823 bp to 253,259 bp) and LS2 (OK638189). The first row shows the bases in the reference sequence, from the second row onwards only the bases that differ from the reference are shown, and the bases that are the same as the reference are indicated by dashes. All insertions are represented by the first two bases and deletions by the letter “D”.

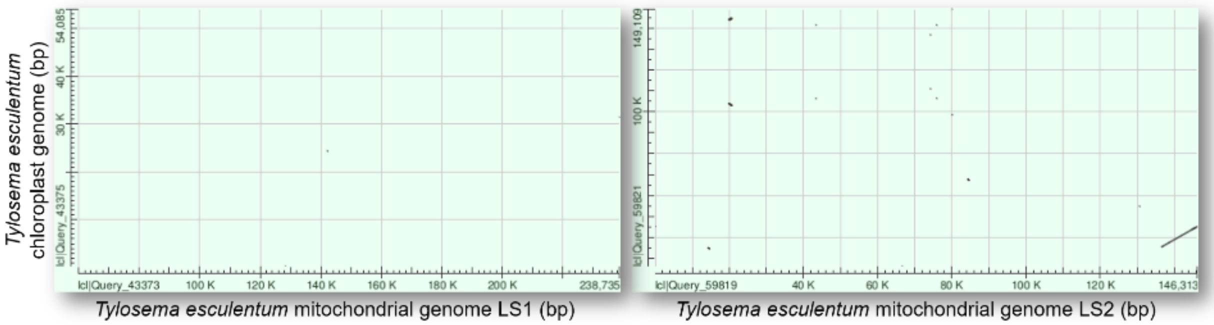


Figure S32. A dot plot view showing the alignment of marama mitochondrial chromosomes to the chloroplast genome. The two reference mitochondrial chromosomes of marama (OK638188 and OK638189) were blasted against the reference chloroplast genome of marama (KX792933.1), respectively in NCBI.

Table S2. List of homologous fragments between the mitochondrial and chloroplast genomes of *Tylosema esculentum*.

| Plastome | | Mitogenome | | | | Alignment | |
|---|---|---|---|---|---|---|---|
| Start | End | Chr | Start | End | Length | %Identity | E-value |
| 260 | 339 | LS1 | 128344 | 128265 | 80 | 98.75 | 4.66E-33 |
| 24312 | 24387 | LS1 | 142158 | 142233 | 76 | 98.684 | 7.80E-31 |
| 26169 | 26237 | LS2 | 66608 | 66676 | 69 | 100 | 1.31E-28 |
| 31434 | 31539 | LS1 | 238735 | 238628 | 108 | 93.519 | 4.63E-38 |
| 34586 | 34943 | LS2 | 14640 | 14299 | 360 | 83.056 | 4.34E-83 |
| 35308 | 45222 | LS2 | 136516 | 146313 | 9926 | 98.146 | 0 |
| 38063 | 38123 | LS2 | 139272 | 139214 | 61 | 88.525 | 6.20E-12 |
| 45221 | 45257 | LS2 | 146312 | 35 | 37 | 100 | 8.02E-11 |
| 54011 | 54085 | LS1 | 59870 | 59796 | 75 | 94.667 | 2.83E-25 |
| 54818 | 54845 | LS2 | 130707 | 130734 | 28 | 100 | 8.08E-06 |
| 67414 | 67689 | LS2 | 84625 | 84353 | 282 | 90.426 | 1.53E-97 |
| 98539 | 98610 | LS2 | 80134 | 80205 | 72 | 97.222 | 6.07E-27 |
| 102961 | 103824 | LS2 | 20757 | 19899 | 891 | 74.074 | 1.21E-83 |
| 106303 | 106383 | LS2 | 43447 | 43375 | 81 | 90.123 | 1.02E-19 |
| 106303 | 106383 | LS2 | 75967 | 75895 | 81 | 90.123 | 1.02E-19 |
| 110921 | 111003 | LS2 | 74229 | 74312 | 84 | 97.619 | 4.66E-33 |
| 136648 | 136730 | LS2 | 74312 | 74229 | 84 | 97.619 | 4.66E-33 |
| 141268 | 141348 | LS2 | 43375 | 43447 | 81 | 90.123 | 1.02E-19 |
| 141268 | 141348 | LS2 | 75895 | 75967 | 81 | 90.123 | 1.02E-19 |
| 143827 | 144690 | LS2 | 19899 | 20757 | 891 | 74.074 | 1.21E-83 |
| 149041 | 149112 | LS2 | 80205 | 80134 | 72 | 97.222 | 6.07E-27 |

Table S3. Potential effect of variations found in the gene sequences on the 9,798 bp cpDNA insertion in the 84 individuals.

| **Position** | **Reference** | **Variation** | **Gene** | **AA Substitution** |
|---|---|---|---|---|
| 35570 | A | C | *psbC* | N44H |
| 35634 | G | T | *psbC* | G65V |
| 35774 | TTTGTGTCT | DEL | *psbC* | FVS112-114Δ |
| 35884 | TTATGTAT | DEL | *psbC* | Y149fs* |
| 36347 | GGACCTACT | DEL | *psbC* | GPT303-305Δ |
| 38753 | CGATCTTC | DEL | *rps14* | G67fs* |
| 38867 | G | T | *rps14* | synonymous |
| 38873 | TTTTTTTGAGGATCGGCGAA | DEL | *rps14* | I23fs* |
| 38893 | T | G | *rps14* | I23L |
| 38922 | CTTTTCTTCT | DEL | *rps14* | E10fs* |
| 39144 | A | G | *psaB* | V711A |
| 39264 | CAATATCCA | DEL | *psaB* | GYW 669-671Δ |
| 39412 | C | A | *psaB* | D622Y |
| 39414 | C | A | *psaB* | R621I |
| 39429 | A | C | *psaB* | L616W |
| 40061 | G | T | *psaB* | D405E |
| 40517 | A | G | *psaB* | synonymous |
| 40544 | A | C | *psaB* | F244L |
| 41030 | A | G | *psaB* | synonymous |
| 41243 | A | G | *psaB* | synonymous |
| 41716 | C | A | *psaA* | D615Y |
| 41718 | G | T | *psaA* | S614Δ* |
| 42060 | GTTGCACCAGGGGCC | DEL | *psaA* | APGAT491-495Δ |
| 42385 | C | T | *psaA* | V392M |
| 42587 | C | T | *psaA* | synonymous |
| 42793 | G | C | *psaA* | Q256E |
| 42817 | CCAAAAGAT | DEL | *psaA* | DLL245-247Δ |
| 42838 | G | T | *psaA* | L241I |
| 43508 | ATCCCTAT | DEL | *psaA* | A15fs* |

Genetic info represented by symbols in the table: AA, amino acids; Δ, deletion; fs, frameshift; *, nonsense mutation.
